# Modeling interpretable correspondence between cell state and perturbation response with CellCap

**DOI:** 10.1101/2024.03.14.585078

**Authors:** Yang Xu, Stephen Fleming, Matthew Tegtmeyer, Steven A. McCarroll, Mehrtash Babadi

## Abstract

Single-cell transcriptomics, in conjunction with genetic and compound perturbations, offers a robust approach for exploring cellular behaviors in diverse contexts. Such experiments allow uncovering cell-state-specific responses to perturbations, a crucial aspect in unraveling the intricate molecular mechanisms governing cellular behavior and potentially discovering novel regulatory pathways and therapeutic targets. However, prevailing computational methods predominantly focus on predicting average cellular responses, disregarding the inherent response heterogeneity associated with cell state diversity. In this study, we present CellCap, a deep generative model designed for the end-to-end analysis of single-cell perturbation experiments. CellCap employs sparse dictionary learning in a latent space to deconstruct cell-state-specific perturbation responses into a set of transcriptional response programs. These programs are then utilized by each perturbation condition and each cell at varying degrees. The incorporation of specific model design choices, such as dot-product cross-attention between cell states and response programs, along with a linearly-decoded latent space, underlay the interpretation power of CellCap. We evaluate CellCap’s model interpretability through multiple simulated scenarios and apply it to two real single-cell perturbation datasets. These datasets feature either heterogeneous cellular populations or a complex experimental setup. Our results demonstrate that CellCap successfully uncovers the relationship between cell state and perturbation response, unveiling novel insights overlooked in previous analyses. The model’s interpretability, coupled with its effectiveness in capturing heterogeneous responses, positions CellCap as a valuable tool for advancing our understanding of cellular behaviors in the context of perturbation experiments.

## 1 Main

High-throughput single-cell RNA sequencing (scRNA-seq) has greatly advanced our understanding of cellular and molecular biology [1–3]. Combining scRNA-seq with perturbation experiments has further expanded our ability to explore the way cells behave in different conditions. Technological innovations like Perturb-seq [4] and CROP-seq [5] use CRISPR to introduce genetic changes or perturb gene expression levels in cells, and they enable systematic screening at large scale. Single-cell perturbation datasets generated using these technologies are promising approaches for discovering comprehensive maps of gene regulatory networks in complex cellular systems [6]. Such an understanding of gene networks could substantially boost drug discovery efforts [7]. As the number of large-scale single-cell perturbation datasets grows, so does the challenge of modeling and interpreting perturbation responses at the single-cell level.

The most straightforward approach to single-cell perturbation data analysis is one aimed at uncovering bulk effects, where expression data from perturbed cells are aggregated and compared to the aggregate of untreated control cells. In practice, addressing cellular heterogeneity (both at the level of cell types and cell states) and batch effects requires complicated data preprocessing steps, including batch effect correction and cell type identification [8]. This rudimentary approach suffers from a number of fundamental shortcomings: (1) The choice on the data preprocessing procedure often lead to changes in biological conclusions; (2) Cell type identification and data stratification often requires imposing arbitrary thresholds (in particular, in experiments dealing with cells on a differentiation trajectory); (3) Bulk analysis testing overlooks potentially valuable information encoded in cell-state heterogeneity, such as cell-state-specific responses; (4) The standard differential expression analysis does not immediately reveal commonalities and contrasts between the mechanism of action of different perturbations, which requires further statistical modeling. It is therefore desirable to address these shortcomings within a robust end-to-end computational framework.

In recent years, several machine learning approaches have made progress toward improving upon simple differential expression testing. The computational framework MIMOSCA assumes an additive model of perturbational responses and analyzes perturbation experiments within a regularized linear regression framework [4]. The computational framework Augur adapts a random forest classifier to prioritize the cell type on which a certain perturbation has primary impact. Once this cell type is identified, a differential expression test can be performed in this cell type context to reveal the perturbation effects [9]. While these approaches recover certain aspects of the underlying biology, it is likely that more complex cellular behaviors are not captured by simple linear models.

Deep learning models have also been explored as a means to uncover the nonlinear complexity in single-cell perturbation data. For example, scGen uses a deep generative model to predict the impact of one perturbation on a new cell population [10], and GEARS takes advantage of prior biological knowledge to model nonlinear gene interactions and nonlinearities in response to multiple perturbations [11]. Though both scGen and GEARS show promise in predicting nonlinear synergistic effects between multiple perturbations, they do not explicitly model the ways in which cell type or cell state generate the nonlinear perturbation response. The Compositional Perturbation Autoencoder (CPA) model, a follow-up to scGen, decomposes the perturbed gene expression profiles into a cell state latent representation in which perturbation, batch, and other effects are all modeled as vector translations [12]. This enables CPA to predict cell-state-specific perturbation responses. Alternatively, PerturbNet encodes perturbation and cell state into two separate latent representations via two different encoding neural networks, and it connects the perturbation representation and cell state representation through a third neural network [13]. These designs allow CPA and PerturbNet to uncover the correspondence between cell state and perturbation response. However, the nonlinear deep neural network decoders involved in the models above do not readily lend themselves to succinct interpretation. Thus, these methods primarily focus on the task of predicting unseen perturbation responses rather than understanding and interpreting the perturbation responses measured by the dataset at hand.

Here we propose CellCap, a linearly-decoded variational autoencoder for modeling single-cell perturbation data. CellCap builds upon the foundation laid by the CPA model but differs in several key ways including its primary objective: while CPA aims to predict the responses of unseen perturbations and in combinations, CellCap instead focuses on dissecting and interpreting cellular responses in terms of a learned sparse dictionary of transcriptional response programs. CellCap combines a nonlinear encoder with a linear decoder. The linear decoder lends the model interpretability by allowing translating all latent space quantities to the gene expression space, while the nonlinear encoder aims to uncover the basal (pre-perturbation) cell state from perturbed cell states. In order to capture the complexity of cell-state-specific responses, CellCap moves the nonlinear computations into the latent space algebra, using multi-head dot-product attention to capture the correspondence between the state of individual cells and their perturbation response. These response amplitudes then act on a sparse dictionary of transcriptional response programs to generate the data. We demonstrate the interpretability and model identifiability of CellCap with both simulated and real single-cell perturbation data.

## 2 Results

### 2.1 CellCap’s key concepts

CellCap is fundamentally a variational autoencoder (VAE) that encodes the observed gene count matrix *x*_*ng*_ into and out of a *K*-dimensional latent space. What distinguishes CellCap from a standard VAE, however, is the additional structure imposed on the making of latent representations. As a first step, the CellCap encoder projects the observed gene count matrix *x*_*ng*_ into a “basal state” 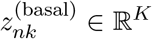 using a multi-layer neural network (Fig. 1a). Here *n* and *g* are cell and gene indices respectively, and *k* is the dimension of the latent space. The latent space modifications arising from fixed effect covariates 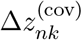 and perturbations 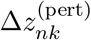 are then added to 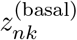 to produce the complete and “recomposed” cell latent representation *z*_*nk*_. The linear fixed effects modeled by 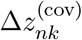 serve to regress out sources of variability that are not the object of study, for example batch or donor identity. The complete latent representation *z*_*nk*_ is then transformed back into gene expression space via a linear decoder and is matched with the observed gene expression matrix *x*_*ng*_ (loss *ℒ*_reconstruction_) [14, 15]. Since the relationship between observed data and the basal state can be highly complex and nonlinear (e.g. the observed data could correspond to treated cells whereas the basal state would correspond to the inferred state of the same cells prior to treatment), we use an expressive deep neural network to amortize the inference of the basal state 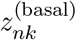 from *x*_*ng*_. At the same time, we use a linear decoder to maintain interpretability of the latent space [16].

**Figure 1.**
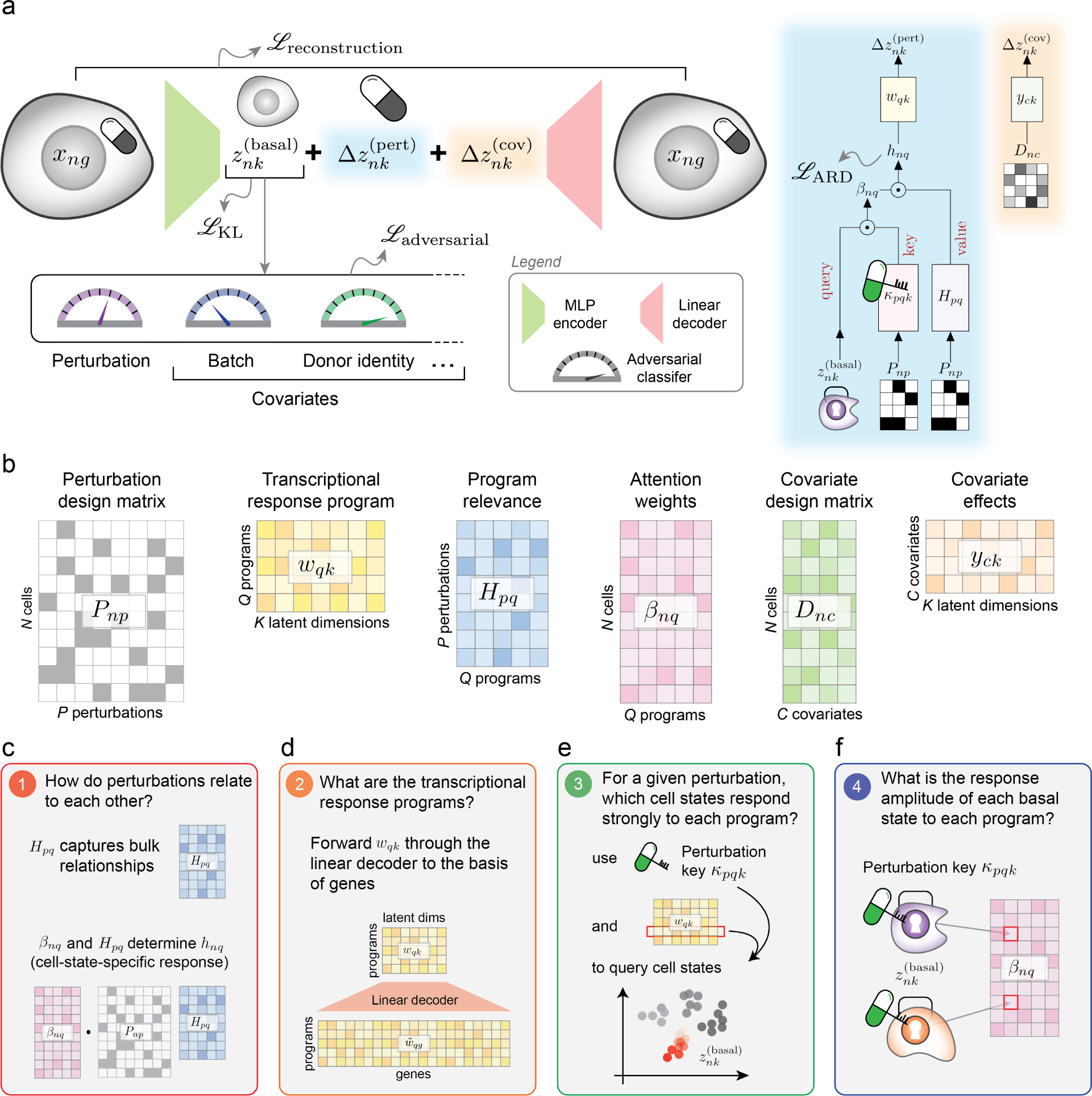
Key concepts underlying the CellCap model and its interpretable workflow. (a) A linearly decoded VAE is the backbone of CellCap, which encodes the perturbed single-cell data into basal state *z*^(basal)^ and reconstructs 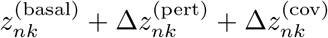 back to the observed gene expression count matrix *x*_*ng*_ via a linear decoder. (b) Graphical presentations of individual components of CellCap. (c-f) Various downstream applications of CellCap. (c) Understanding the relationship between bulk and cell-state-specific perturbation effects; (d) Interpretability of the transcriptional response programs with a linear decoder; (e) Uncovering characteristics of cell states that respond to each transcriptional program; (f) Uncovering cell-state-dependent response amplitude to each response program.

The “basal state” is a concept proposed in the CPA model by Lotfollahi *et al*. [12] and is understood as an intermediate latent space vector that captures only the intrinsic and unmodeled cell state variation. Throughout this manuscript, the terms “cell state” and “basal state” are used interchangeably. Importantly, 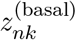 ought not to contain information related to perturbation and other known covariates such as batch or donor identity. To obtain such a basal state representation, we simultaneously train the encoder network and adversarial classifier networks to strip perturbation and known covariate information out of the basal latent space, similar to the approach taken by the CPA model (loss *ℒ*_adversarial_).

We formulate modeling the effect of perturbations as a “dictionary learning” problem. We assume that the applied perturbations can induce up to *Q* transcriptional “response programs”. The response programs can be collected as a matrix *w*_*qk*_ *∈* (−1, 1) interval, with *Q* rows, with each row representing a distinct response program. All cells share statistical power to estimate *w*_*qk*_, though individual programs can be used by different cells and perturbations with different amplitudes (Fig. 1a). Explicitly, we assume 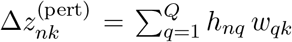, where the *h*_*nq*_ represents the to-be-inferred usage amplitude of response program *q* by cell *n*.

Our use of low-complexity building blocks for the sake of interpretability, namely a linearly-decoded latent space and dictionary learning, necessitates invoking an element of nonlinearity to allow modeling complex data distributions beyond the reach of linear models. In our formulation, we introduce this nonlinearity in the computation of the variational posterior distribution of *h*_*nq*_, which is one of the key methodological contributions of our work. We construct *h*_*nq*_ in the form of “scaled dot-product attention” [17] between the basal cell state 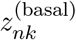 and perturbation design matrix *P*_*np*_ ∈ {0, 1}. The perturbations are associated with a set of learned “perturbation key” vectors ***κ***_*pq*_ (or equivalently, a 3-dimensional tensor *κ*_*pqk*_). The key vectors, which live in the same *K*-dimensional space as the basal states (see Eqn. 3), determine the amplitude of cell-state-specific response program *q* in cell *n* as 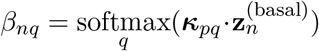. Illustrated in Fig. 1a, the entries of *β*_*nq*_ denote the normalized attention weights between cells and response programs. In the terminology of scaled dot-product attention, the the basal state serves as “query”, the perturbation keys serve as “key”, and finally the “value” is *v*_*nq*_ = Σ_*p*_ *H*_*pq*_ *P*_*np*_, which is intuitively the linear action of *H*_*pq*_ on the *P*_*np*_. Here, *H*_*pq*_ is another learnable weight matrix, with the interpretation of the usage of response program *q* by perturbation *p* at an aggregate level. Ultimately, the program usage is given as *h*_*nq*_ = *β*_*nq*_ *v*_*nq*_ and can be interpreted as the cell-state-specific amplitude of response program *q* in cell *n* after being treated with perturbation *p*. In practice, as in Ref. [17], we also found it beneficial to leverage multiple attention heads to distribute the inference of response program usages to specialized attention heads (see Methods).

We remark that the learned dictionary of response programs *w*_*qk*_ are reused across perturbations, leading to improved interpretability of the output and an understanding of the relatedness between perturbations. CellCap uses sparse Bayesian learning (SBL), in the form of “automatic relevance determination” [18], as a mechanism to learn as few response programs (*<*= *Q*) as are necessary to explain the data (loss ℒ_ARD_). The total loss function of CellCap balances this learning objective with the reconstruction loss and the adversarial loss (Fig. 1a). A formal definition of the generative and inference process is provided in the Methods section, as well as hyperparameter setup for balancing these learning objectives.

The CellCap model lends itself to interpretability by design (Fig. 1b). Here, we outline a few biological questions that can be answered using CellCap. First, CellCap reveals whether different perturbations elicit similar cellular responses (Fig. 1c). General relationships between perturbations are captured by *H*_*pq*_, the usage of each learned response program by each perturbation. CellCap enables researchers to ask the same question under a specific cell-state context by examining *h*_*nq*_, which is a combination of *H*_*pq*_ and *β*_*nq*_, the cell-state-dependent attention outputs that describe the response amplitude of each response program in each perturbed cell. This enables a finer-grained understanding of relationships among perturbations while going beyond bulk effects and leveraging the single-cell resolution of the data. Next, CellCap defines the transcriptional response programs activated in a single-cell perturbation experiment (Fig. 1d). Instead of an overall effect for each perturbation, CellCap discovers individual transcriptional response programs in the form of *w*_*qk*_. Since these programs live in the same latent space as cell states, we can use the linear decoder to translate these latent-space response programs to gene expression space, where each program can be understood in terms of gene expression patterns.

The key advantage of CellCap is its ability to model the correspondence between basal cell states and perturbation responses (Fig. 1e). This correspondence is critical for understanding why cells may respond to a perturbation in a particular way. For example, the activation of cell-state-specific gene expression programs, which could include cell maturity or cell cycle phase among others, may make a specific cell population uniquely vulnerable to a perturbation and lead them to respond with a specific transcriptional activation. This correspondence can be uncovered by using the perturbation key *κ*_*pqk*_ of perturbation *p* in the context of response program *q*, to query which basal state 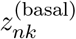 is relevant (see Methods). Finally, the attention weights *β*_*nq*_ indicate how the amplitude of response program *q* varies across different basal cell states (Fig. 1f).

### 2.2 CellCap captures cell-state dependent responses in simulated data

To explore different aspects of the behavior of CellCap in a controlled fashion, we generate and study three simulated scenarios that contain two perturbations in each (Fig. 2a). We assume that the basal cell states co-vary with a hypothetical “pseudotime” to serve as a simplified model of cell state trajectories. We used PROSSTT [19] to generate such continuous cell states, and each cell was assigned a pseudotime value (see Supplementary Section S.1). In simulated scenario (1), each perturbation induces a single unique transcriptional response program, with the amplitude of the response being proportional to the cell state “pseudotime”. In simulated scenario (2), the two perturbations induce a single shared response program, though the cells have a stronger response to one perturbation than the other. Again, the cell state pseudotime modulates the amplitude of the responses similarly for both perturbations. In simulated scenario (3), each perturbation induces one shared response and one unique response. In this case, the shared response amplitude is correlated with the cell state pseudotime, and the unique responses are anti-correlated with the pseudotime.

**Figure 2.**
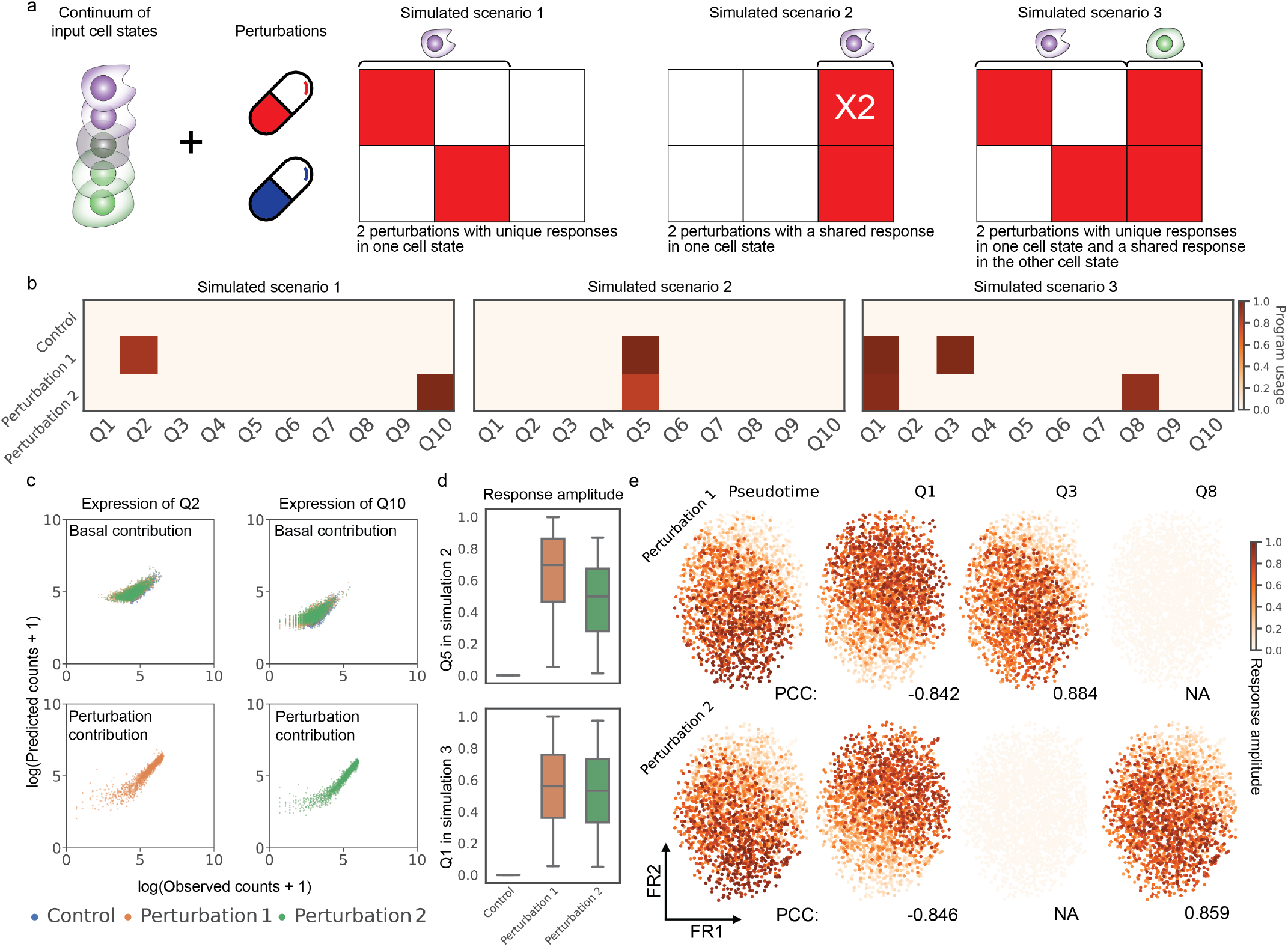
Exploring CellCap’s model identifiability using simulated data. (a) Graphical setup of 3 simulated scenarios. (b) Usage of each transcriptional response program discovered by CellCap (*h*_*nq*_ averaged over cells). Automatic relevance determination effectively turns off response programs that are not necessary to explain the data, in agreement with the ground truth. (c) The linear decoder is used to directly interpret the learned basal state 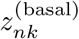 and the learned perturbation 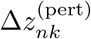. Each dot represents the sum of all relevant genes in one cell for the indicated response program, and cells are colored by perturbation condition. CellCap learns a basal state where perturbations are indistinguishable (top panels), while learning response programs whose expression correlates with ground truth (bottom panels). (d) Amplitudes *h*_*nq*_ of the shared response programs (Q5 in scenario (2) and Q1 in scenario (3)) in each perturbation condition are presented with boxplots. (e) Visualization of per-cell response amplitudes *h*_*nq*_ for the 3 learned response programs in scenario (3). The leftmost Fruchterman-Reingold (FR) plots are colored by the ground truth cell state pseudotime, whereas *h*_*nq*_ is used to color the FR plots shown in rightmost three columns. The pseudotime hidden variable is the ground truth covariate of the cell state response amplitude (see Supplementary Sec. S.1 for details). Unique (shared) response programs are expected to be correlated (anti-correlated) with the pseudotime.

In scenario (1), we expect that CellCap should identify two response programs and that each response should be specific to one perturbation. Indeed, only two programs stand out after model training (Fig. 2b). CellCap also ascertains that there is only one relevant response program in scenario (2) and three in scenario (3). In scenario (2), CellCap identifies response program *Q5* as a shared program induced by both perturbations (Fig. 2b). In scenario (3), CellCap identifies response program *Q1* as the shared program, and response programs *Q3* and *Q8* as the programs specific to perturbation 1 and perturbation 2, respectively (Fig. 2b). We also note that in all cases, we have allowed CellCap to learn and use up to 10 response programs. However, the ARD sparsity-inducing mechanism in CellCap correctly “turns off” the unnecessary transcriptional programs.

Next, we examined whether CellCap accurately decomposed the observed gene expression into basal state (cell state) and response program in the latent space. To this end, we decoded the learned **z**^(basal)^ to obtain the predicted basal gene expression matrix. Our expectation is that the results would be the same across all three conditions, and only exhibit the variation related to pseudotime and not the specific perturbations. We further decoded the complete latent state vector **z** = **z**^(basal)^ + Δ**z**^(pert)^ to predict a full gene expression matrix. We subtracted the predicted basal gene expression matrix from the full gene expression matrix to obtain the predicted perturbation-induced gene expression. As expected, we found the basal state expression to be indistinguishable among control and perturbed cells, and the expression of response programs to be perturbation-specific (Fig. 2c). Additionally, CellCap successfully decomposed the observed gene expression into basal state and response programs in scenarios (2) and (3) (Supplementary Fig. S1). Meanwhile, we found that CellCap accurately learns response programs that match the simulated ground truth (Supplementary Fig. S2).

As mentioned earlier, the two perturbations induce a shared response program in both scenarios (2) and (3), with an unequal response amplitude in scenario (2) and an equal response amplitude in scenario (3). We found that CellCap correctly identified this difference in scenario (2) and reported similar response amplitudes in scenario (3) (Fig. 2d). In scenario (3), we have set up a situation in which the shared response program’s amplitude should be negatively correlated with the basal state “pseudotime”, in contrast to the unique response programs which are positively correlated with basal state pseudotime. The response amplitude *h*_*nq*_ reported by CellCap in scenario (3) was highly correlated with ground truth (Pearson correlation coefficient 0.884 for Q3 in perturbation 1 and 0.859 for Q8 in perturbation 2). As for the shared program, CellCap recovered a shared response program Q1 that captures the negative correlation with ground truth basal state, with Pearson correlation coefficient -0.842 in perturbation 1 and -0.846 in perturbation 2 respectively (Fig. 2e). In summary: (1) CellCap was able to identify shared and unique programs for two perturbations; (2) CellCap could distinguish different response amplitudes; and (3) CellCap learned the correct correspondence between basal cell state and perturbation response.

### 2.3 CellCap reveals heterogeneous responses in pathogen-exposed human monocytes

We next sought to demonstrate the utility and interpretability of CellCap in real single-cell perturbation data. To this end, we used CellCap to reanalyze a previously published pathogen-exposed human peripheral blood mononuclear cell scRNA-seq dataset. In the original report, Oelen *et al*. identified differentially expressed (DE) genes by comparing each treatment condition against untreated control for every major cell type [20]. They observed the largest number of DE genes in monocytes across different pathogen exposure conditions, and they concluded that monocytes are the cell type with the strongest response to pathogens. Another highlight of their analysis is that the interferon signaling pathway is specifically enriched at 3 hours post-exposure in monocytes and that this response pathway is common to all three pathogens. The original analysis did not leverage the single-cell resolution of the dataset for studying the complex and heterogeneous cell-state-dependent responses to pathogen exposure. Here, we show that CellCap can be utilized to uncover such novel insights about cellular responses at an increased granularity. The steps outlined below demonstrate the overall workflow other practitioners could use to leverage CellCap in studying their single-cell perturbation experiments.

First, we limited the scope of our analysis to the monocytes in the dataset, since this was the cell type observed to exhibit the strongest responses in the original study. We noticed that CellCap’s encoding of cell states from different conditions into the shared basal state 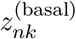 results in excellent mixing between the pathogen-exposed group and the untreated (UT) group (Fig. 3a). We recall that the basal state representation should preserve only intrinsic cell state variations and not the perturbations themselves. Examining the learned usage of each response program by each perturbation condition, we were able to paint an overall picture of the relationship between the 6 treatment conditions by performing principal component analysis (PCA) on the perturbation signatures, i.e. the rows of *H*_*pq*_. In particular, PC1 separated the conditions by time post exposure: all treatments 3 hours (3h) post pathogen exposure – *C. albicans* (CA), *M. tuberculosis* (MTB), and *P. aeruginosa* (PA) – were closely grouped in the PC space, while all 3 treatments 24 hours (24h) post pathogen exposure were located on the right side of the plot (Fig. 3b). Importantly, this indicates that the top PC of response program usage by perturbation is the time post exposure rather than the pathogen itself. Of note, pathogen exposure with PA was distinct from the other two pathogens at the 24h timepoint. We further examined *h*_*nq*_, the usage of each response program by individual cells, and averaged over the top 90 percent of cells with the highest responses to the treatment conditions (Fig. 3c). We observed that all 3 treatments 3h post pathogen exposure primarily induce two programs (Q6 and Q9) but that all treatments 24h post pathogen exposure induce more diverse response programs, including Q3, Q4, Q7, Q8, and Q10 (Fig. 3c). We also note that Q2 and Q5 were not used, i.e. they were turned off during the course of model training, suggesting that the experiment can be succinctly interpreted using 8 response programs.

**Figure 3.**
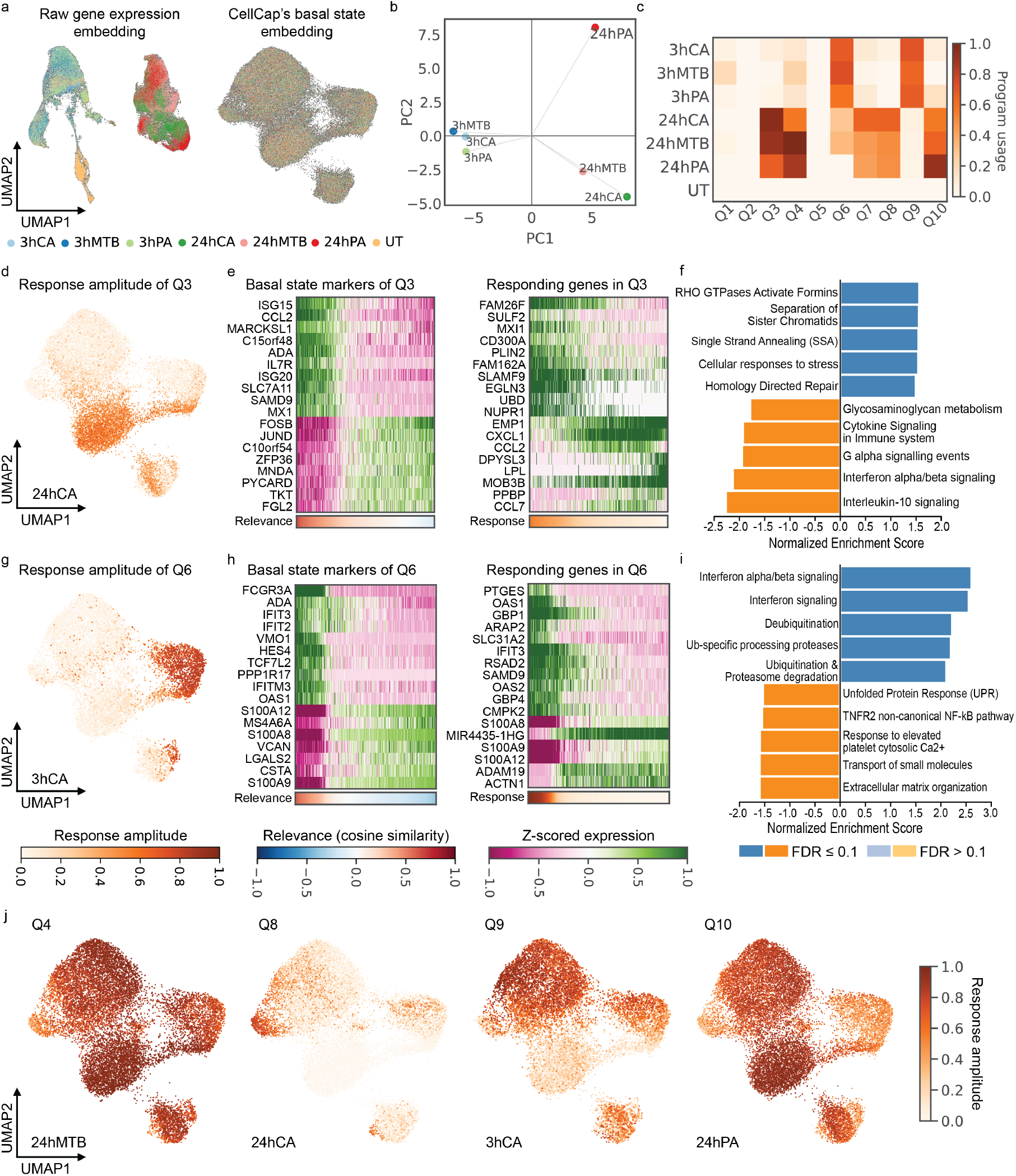
CellCap reveals heterogeneous responses in pathogen-exposed human monocytes. (a) The left and right UMAPs show the embedding of the raw gene expression and CellCap-inferred corresponding basal states, respectively. CellCap learns a basal latent space in which cells from all perturbation conditions are well-mixed. (b) General relationship between 6 pathogen-exposure conditions, visualizing *H*_*pq*_ using principal component analysis. (c) Program usages in each condition, *h*_*nq*_, summed over responding cells in each perturbation group (see Supplementary Section S.2). (d) Per-cell response amplitudes *h*_*nq*_ of program Q3 in the 24hCA perturbation condition. (e) Top basal marker genes and responding genes in Q3. Top up- and down-regulated genes are shown to highlight expression patterns on both extremes. (f) GSEA for response program Q3. Top 5 positive and top 5 negative pathways are shown. (g) Per-cell response amplitudes of program Q6 in the 3hCA perturbation condition. (h) Top basal marker genes and responding genes in Q6. Top up- and down-regulated genes are shown to highlight expression patterns on both extremes. (i) GSEA for response program Q6. (j) Other major cell-state-specific response patterns identified by CellCap. Further investigation of these patterns is shown in Supplementary Figs. S6-S10. Abbreviations: [3hCA = 3 hours post *C. albicans* exposure, 24hCA = 24 hours post *C. albicans* exposure, GSEA = gene set enrichment analysis]

Having obtained a global understanding of both the heterogeneity of basal cell states within the monocyte population (Fig. 3a) and the relationships between different treatment conditions (Fig. 3b and c), we next examined the way in which transcriptional response programs are activated in different basal states of monocytes. We started with program Q3, which is common to all 3 treatments 24h post pathogen exposure. Response amplitudes *h*_*nq*_ of individual cells 24 hours after CA exposure show that the response program Q3 is enriched in a certain sub-population of monocytes (Fig. 3d). We then examined the characteristics of cell states within this sub-population of monocytes. To this end, we used the perturbation key *κ*_*pqk*_ for response program Q3 at 24h post CA exposure to query the basal state 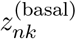 of the UT group. We assigned to each untreated cell a relevance score (computed using cosine similarity) to represent the likelihood of this cell to respond with program Q3 in the 24h post CA exposure condition given its basal state (Supplementary Fig. 3a). We used this relevance score to identify the corresponding basal expression program enriched in cells with a high relevance score (see Methods).

The basal expression program included top upregulated genes *ISG15, CCL2, ISG20*, and *IL7R* (Fig. 3e left panel). We then identified the most highly responding genes in response program Q3 (Fig. 3e right panel). We noted a striking correspondence between basal expression of specific genes (Fig. 3e left) and the activation of Q3 (Fig. 3e right) in response to CA exposure at the 24h timepoint. Gene set enrichment analysis (GSEA) indicates that the response program Q3 is enriched for pathways involved in Rho GTPase activation and DNA replication with FDR ≤ 0.1 (Fig. 3f).

Using the same approach, we identified the basal expression program which results in a high response of program Q6 in the 3h post CA exposure condition (Fig. 3g-i and Supplementary Fig. S3b). We also examined the averaged and individual expression patterns of responding genes in Q3 and Q6 to confirm substantial perturbation changes from the untreated group (Supplementary Figs. S4 and S5). The primary observation is that treatments 3h post exposure induce a higher response of the interferon signaling pathway, and this response is enriched in non-classical monocytes with marker genes *FCGR3A* and *HES4* identified in basal cell state expression. This result is consistent with the finding in the original report, which used a multi-step analysis (clustering, cell state annotation, and case-control comparison) [20]. CellCap not only automates this discovery workflow within a unified end-to-end model, it also uncovers novel response patterns which we briefly outline below.

Additional response patterns identified by CellCap are shown in Fig. 3j and Supplementary Fig. S6. With these patterns in hand, we identified all major basal expression programs and performed soft annotation to assign cellular identities to different sub-populations of monocytes in the basal state without clustering (Supplementary Fig. S7). One salient example is the enrichment of Q8 in a small population of monocytes at 24h post CA exposure. We identified its corresponding basal expression program using the same approach as above (Supplementary Fig. S8). As mentioned earlier, PA pathogen exposure is distinct from the other two pathogens at the 24h timepoint. A possible explanation for this difference could be the enrichment of response program Q10 in one particular sub-population. Q3 is enriched in the same sub-population of monocytes in the all 24h post exposure conditions. However, cells in this sub-population show a higher response of Q10 to 24h PA than to the other two pathogens at 24h. Averaged expression patterns of perturbed genes in Q3 and Q10 indicate that Q3 is a shared program across the 3 pathogens but Q10 is specific to PA exposure (Supplementary Fig. S9). In Q10, we identified macrophage markers like *APOE, APOC1*, and *RNASE1* (Supplementary Fig. S10). This result suggests a possibility that cells in this sub-population of monocytes (with basal marker genes *ISG15, CCL2, ISG20*, and *IL7R*) would gain the potential to differentiate into macrophages after 24 hour exposure to PA. We remark that this nuanced finding enabled by CellCap was not part of the original report by Oelen *et al*. [20].

### 2.4 CellCap captures complex patterns in large-scale genetic perturbation data

We next used CellCap to analyze a single-cell Perturb-seq dataset by Norman *et al*. [21]. The study used CRISPR activation to over-express transcription factors in the K562 cell line at large scale, including 105 single-target perturbations and 131 pairwise combinatorial perturbations. In CellCap, this experimental design can be readily encoded as a design matrix *P*_*np*_ ∈ {0, 1} (with the *p* dimension of size 105), where Σ_*p*_*P*_*np*_ = 1 for the cells receiving single-target perturbations, and Σ_*p*_ *P*_*np*_ = 2 for the cells receiving double-target perturbations. We initially set up a maximum of *Q* = 50 response programs for CellCap to learn. Following model training, we observed that a majority of programs were turned off, leaving only 10 programs shared by the 236 perturbations (Supplementary Fig. S11).

We obtained averaged perturbation response signatures, defined as the learned usage of each response program averaged over responding cells (see Supplementary Section S.2) stratified by each of the 236 perturbation conditions. We performed UMAP dimensionality reduction on the obtained average perturbation signatures for visualization, and we clustered the signatures (Fig. 4a). This clustering largely agrees with the findings in the original study by Norman *et al*. [21] and a reanalysis done by Roohani *et al*. [11]. Norman *et al*. clustered the perturbations using a pseudo-bulk approach, while Roohani *et al*. used their proposed deep learning method GEARS.

**Figure 4.**
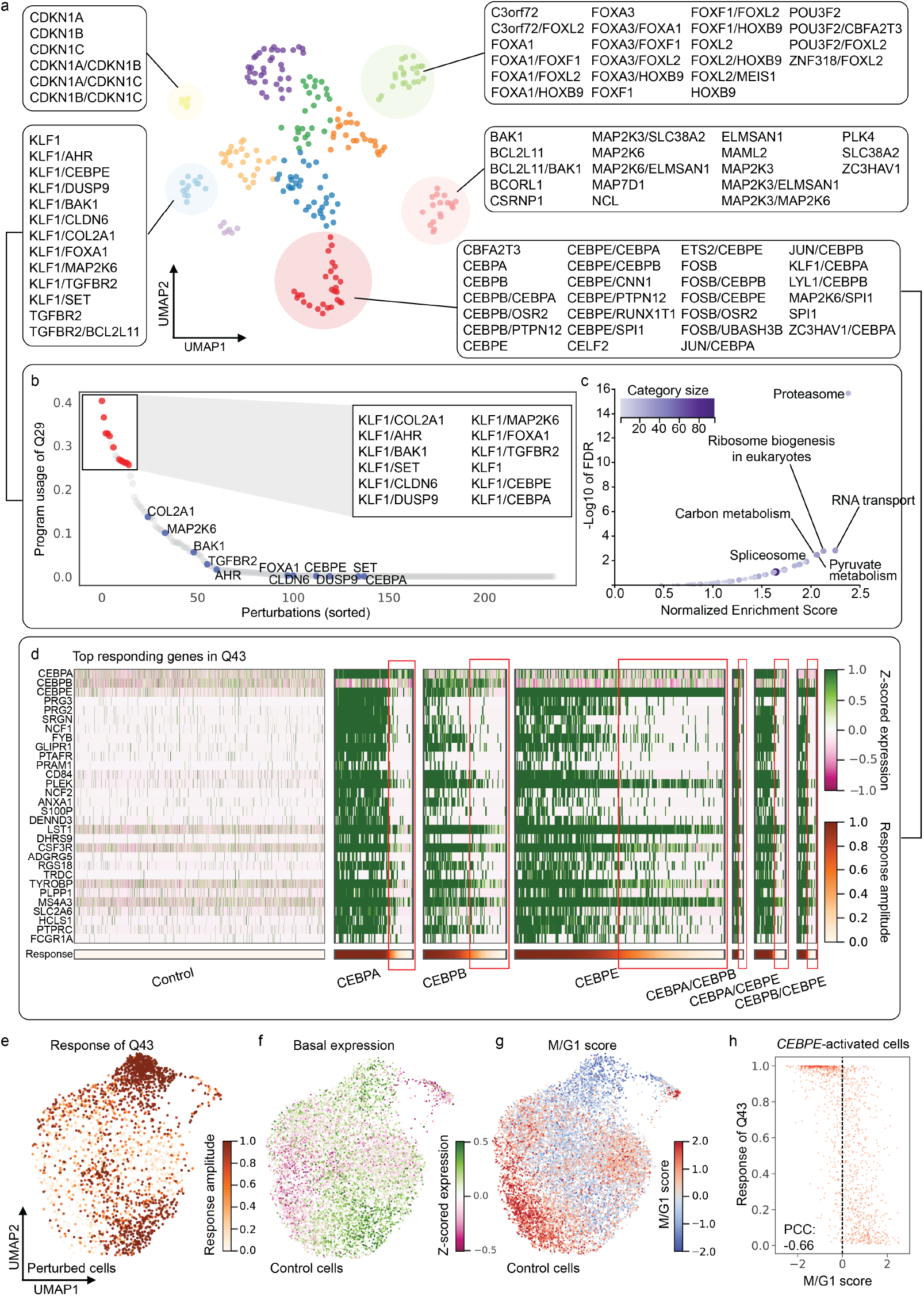
CellCap captures complex relations between genetic perturbations in large screens (data from Norman et al. [21]). (a) UMAP visualization of the relatedness of 236 perturbations obtained from the program usages inferred by CellCap (Supplementary Section S.2). Perturbations are colored by K-means clustering, highlighting 5 perturbation clusters. (b) Perturbations ranked by usage of the response program Q29. Perturbations involving *KLF1* activation are colored in red while others are colored in grey. Blue dots highlight the single-target perturbations (that lack *KLF1* activation) corresponding to those red perturbations. (c) GSEA results showing significant KEGG pathways related to response program Q29. KEGG pathways with FDR ≤ 0.1 are highlighted. Dot size and color are proportional to the size of the gene set. (d) Expression of top responding genes in program Q43. For each perturbation, cells are ordered from the highest Q43 response amplitude to the lowest. (e) Per-cell usage *h*_*nq*_ of the Q43 response program are shown on the basal state UMAP for all the perturbed cells from panel (d). (f) The basal state UMAP of control cells showing the average z-scored expression of the top basal state marker genes that correspond to program Q43. (g) Same basal state UMAP showing M/G1 score for control cells. (h) Quantitative comparison of M/G1 score and Q43 response amplitude in *CEBPE* -activated cells reveals a strong correlation.

We focused on 4 of these perturbation clusters for downstream investigation. Ranking the perturbations by their usage of response program Q29, we found that the top ranked perturbations primarily involve the activation of *KLF1* as well as *KLF1* -included combinatorial perturbations (Fig. 4b). We noticed that perturbations like *AHR, BAK1, DUSP9, SET* are ranked lower than their combinational perturbatins with *KLF1* in response program Q29 (Fig. 4b). We wondered whether response program Q29 is primarily driven by activation of *KLF1*. We examined the expression of top responding genes in Q29 in perturbation conditions involving the activation of *KLF1, AHR, BAK1, DUSP9, SET*, and their pairwise combinations *KLF1/AHR, KLF1/BAK1, KLF1/DUSP9* and *KLF1/SET*. We found that the top responding genes in response program Q29 are primarily expressed in *KLF1* -activated perturbations but not in *AHR, BAK1, DUSP9* -or *SET* -activated perturbations (Supplementary Fig. S12). This indicates that response program Q29 is mainly caused by activation of *KLF1*, with no or little dependence on the presence of *AHR, BAK1, DUSP9* or *SET* activation.

Activation of *KLF1* has been shown to promote erythropoiesis [22, 23]. However, the analyses in both Norman *et al*. [21] and Roohani *et al*. [11] categorized perturbations with activation of *KLF1* as “pro-growth”. We performed GSEA and found that response program Q29 might be related to metabolic processes or proteasome activation (Fig. 4c). Meanwhile, CellCap identified lymphocytes marker genes in response program Q29. These lymphocytes markers include *CD3G, CD80*, and *IL2RG* (Supplementary Fig. S12). Response program Q29 also includes other non-specific markers. For example, *TRABD2A* and *NMU* also have high expression in lympho-cytess. Based on this, we hypothesize that *KLF1* activation in K562 cells could serve to enhance lymphocytes identity.

Both Norman *et al*. and Roohani *et al*. identified the group of perturbations that includes *CNN1, CBL*, and *UBASH3B* as “erythroid”. Consistent with their analyses, CellCap also identified that erythroid markers like *HBE1, HBG1*, and *HBG2* are highly ranked in the response program Q7 shared by these perturbations (Supplementary Fig. S13). Interestingly, CellCap also revealed *KLF1* as one of top responding genes in response program Q7, suggesting *KLF1* may promote erythropoiesis indeed [22, 23]. Although both Norman *et al*. and Roohani *et al*. labeled the group of perturbations that includes *FOXA1, FOXA3*, and *FOXL2* as “pioneer factors”, a detailed description of shared perturbation responses in this group is missing. Here, CellCap identified that this group of perturbations shares response program Q22, which includes top responding genes like *LYZ* and *ID3* (Supplementary Fig. S14).

We next focused on response program Q43, which is most strongly induced by activation of *CEBPA, CEBPB, CEBPE*, and their combinatorial perturbations. Both Norman *et al*. and Roohani *et al*. classified this group as granulocytes [11, 21]. CellCap identified neutrophil markers in Q43, including *LST1* and *CSF3R* (Fig. 4d), showing great agreement with previous analyses. Weinberger *et al*. also reanalyzed this Perturb-seq dataset with a primary focus on this group of perturbations [24]. By visualizing a few neutrophil marker genes, they confirmed that only a certain population of cells in which *CEBEB* and/or *SPI* are activated would differentiate towards neutrophils. However, Weinberger *et al*. did not highlight the heterogeneity in response amplitudes at the single-cell level in the *CEBPB* and *SPI* -activated perturbations. CellCap reveals that this heterogeneity in per-cell response magnitude occurs in all perturbations involving *CEBPA, CEBPB, CEBPE*, and their combinations (Fig. 4d: highlighted with red boxes). Expression levels of genes *CEBPA, CEBPB*, and *CEBPE* (the target genes) suggest that this heterogeneity is not likely due to a failure to activate the target gene in these cells (Fig. 4d: top 3 rows). For example, the gene *CEBPE* is robustly activated in all perturbed cells in the *CEBPE* perturbation group, even though the cells highlighted by the red box show a much lower level of the Q43 response program.

This motivated us to examine the basal cell state to understand if some intrinsic, pre-existing cellular variation would explain this phenomenon. We computed a cosine similarity between the perturbation key *κ*_*pqk*_ of *CEBPA, CEBPB, CEBPE* or *SPI1* and the basal state 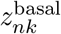 of control cells, and we identified a basal program related to response program Q43. We compared the pattern of response amplitudes of program Q43 in perturbed cells (Fig. 4e) with the average expression of the related basal program in control cells (Fig. 4f), and we confirm that the patterns match. Regarding this related basal program, the unperturbed K562 cells with high cosine similarity to all 4 perturbations already present low expression of the granulocytes marker genes *CSF3R, LST1, LGALS1*, and *APOC1* (Supplementary Fig. S15). This indicates the existence of or ongoing differentiation towards granulocyte lineage in unperturbed K562 cells. CellCap finds that cells which display some granulocyte markers in the basal state will continue on a path toward granulocyte differentiation as a response to *CEBP\** activation.

CellCap also indicates that cell cycle stage is anti-correlated with Q43 response. We visualized cell cycle patterns in control cells (Fig. 4g and Supplementary Fig. S16) and confirmed that a per-cell M/G1 score shows strong anti-correlation with the magnitude of Q43 response (Fig. 4h; with additional perturbations shown in Supplementary Fig. S17). CellCap pinpoints the cell cycle genes that are anti-correlated with program Q43. This anti-correlation is especially strong for *CENPA, CENPE* and *CENPF*, which are known marker genes for the M stage. Cells with lower Q43 response amplitudes tend to have higher expressions of *CENPA, CENPE* and *CENPF* (Supplementary Fig. S18). Weinberger *et al*. also found that cell cycle is a shared variation across conditions [24]; however, their analysis did not uncover the anti-correlation between M/G1 score and the transcriptional response to activation of *CEBPA, CEBPB*, and/or *CEBPE*. CellCap’s nuanced findings suggest a possibility that cells that already show signs of differentiation towards granulocytes and are not in G1/M phase would have higher response to activation of *CEBPA, CEBPB*, and *CEBPE*.

## 3 Discussion

In this study, we developed the computational method CellCap for end-to-end analysis of single-cell perturbation experiments. CellCap models an interpretable correspondence between cell states and perturbation responses and enables multiple downstream applications. We demonstrated the utility of CellCap and the consistency of its findings using both simulated and real single-cell perturbation datasets. Our reanalysis of pathogen-exposed human monocytes data from Oelen *et al*. [20] uncovered novel cell-state-dependent responses following pathogen exposure. In particular, we found a sub-population of monocytes that exhibit macrophage differentiation potential 24 hours after *P. aeruginosa* exposure, a finding that was overlooked in the original study. Our reanalysis of genetic perturbation data collected by Norman *et al*. [21] demonstrated agreement with previous analyses while also providing further insights into the ways in which the effects of genetic perturbations can be nuanced in relation to cell states.

The methodological innovations of CellCap are to be understood in relation to the existing methods developed in this domain which put different degrees of emphasis on interpretability and prediction accuracy. One of the key concepts underlying CellCap is that each cell is endowed with “basal state”, an inferred pre-perturbation transcriptional state that encodes intrinsic cell state variation. While we have borrowed this idea from the CPA model [12], we emphasize that the end goal of CellCap is interpreting and decomposing single-cell perturbation responses into a dictionary of response programs, in contrast to predicting unseen perturbation effects accurately. Even though both CPA and CellCap aim to model the correspondence between basal cell state and perturbation response, the interpretive advantage of the CellCap model lies in the additional structure imposed on the latent space operations. CellCap learns this correspondence explicitly in the form of attention weights that signify the coupling between basal state sub-populations, learned response programs, and the usages of these response programs within each perturbation condition. While CPA has the potential to exhibit greater accuracy at predicting perturbation effects in unseen conditions (using high-dimensional latent spaces and nonlinear decoders), CellCap’s utility lies in the complementary role it plays in dissecting perturbation experiments and transforming observed cellular responses into biological insights.

The computational tool contrastiveVI [24] encodes single-cell perturbation data into independent “shared” and “salient” latent spaces, and is another valuable method for dissecting single-cell perturbation data. The concepts of “shared” and “salient” representations in contrastiveVI have semblance to the separable basal state and perturbation representations in CellCap and CPA. One key difference is that contrastiveVI does not model the correspondence between the “shared” and “salient” components (i.e. the equivalent of CellCap’s attention structure), and as such, the model does not explicitly uncover the relationship between perturbation response and cell states. This difference was demonstrated by the results highlighted in Section 2.4 in which we reanalyzed the Perturb-seq data from Norman *et al*. [21]. CellCap was able to uncover previously unnoticed patterns in heterogeneous perturbation responses caused by interpretable differences in cellular basal state – in this case, pre-existence of granulocytes lineage and cell cycle phase. Learning the correspondence between basal cell state and perturbation response not only enhances interpretability, but also gives CellCap greater statistical power for response program discovery as opposed to more direct differential expression testing methods, which find transcriptional responses that are averaged over responding and non-responding cells.

Unsupervised Bayesian models such as CellCap, though highly effective in extracting insights from new datasets, come with certain trade-offs and practical considerations. One of these challenges involves determining appropriate hyperparameter values. For instance, the line between a “shared” and “unique” response program can be hard to draw in certain cases. There is a necessary trade-off between the model’s ability to explain the data *accurately* and its ability to explain the data *concisely*. Shared response programs lend interpretability to the output, but a model that finds additional unique responses for each perturbation can provide a better fit to the data. We make this trade-off transparent and explicit in CellCap by means of a hyperparameter that controls the weight given to automatic relevance determination in the optimization objective function. A larger value will encourage CellCap to explain the data using fewer response programs, whereas a smaller value will encourage the model to reconstruct the data more accurately at the expense of learning more response programs. This trade-off is discussed in more detail in Methods 4.1.6. Researchers are encouraged to carefully scrutinize the hyperparameters for each dataset, making thoughtful choices to align with other relevant factors and the overarching objectives of the study. It is essential to learn a biologically meaningful basal state in both CellCap and CPA, akin to the”shared” representation in contrastiveVI. However, there are scenarios where inferring the basal state could prove challenging without gathering time courses of higher resolution. For instance, in a diving cell culture, perturbations could lead to the emergence of a new cell state, coupled with the elimination of specific cell sub-populations and/or a near-total loss of the original cell state post-perturbation. Along these lines, Weinberger *et al*. have commented that the results produced by constrativeVI can be misleading if the “shared” background variations are not present in perturbed cells [24]. Likewise, CellCap is likely to fail to correctly project perturbed cells back to the pre-perturbation basal state, one that matches the distribution of control cells, if the perturbed cells have undergone significant reprogramming or passages. The evidence for such a failure mode would be (1) obtaining basal state distributions that do not match the state of unperturbed control cells, combined with (2) the use of excessively large values for the hyperparameter *γ* that controls the alignment between basal state distributions across different conditions (see 4.1.6 in Methods). In such a case, downstream analyses and other CellCap inferences are questionable, and obtaining a finer time course could provide the most robust remedy.

Careful consideration is essential when examining perturbation responses in diverse cell populations. It is generally expected that the greater the heterogeneity within the cell population, the more varied the perturbation responses will be. However, when conducting experiments with multiple cell types and seeking to understand and analyze the impact of perturbations on each cell type with the same level of detail, challenges may arise in choosing a single set of hyperparameter values. For instance, cell types that exhibit strong responses may benefit from stronger ARD regularization to reduce the number of learned programs, but this decision may sacrifice granularity in modeling the effects of perturbations in less responsive cell types. In our study, we specifically trained CellCap on datasets featuring a single cell type, with a focus on addressing variations at the fine-grained resolution of cell state. It is important to note that this limitation is not unique to our methodology. For instance, GEARS must also be trained on a single cell type due to potential variations in interactions between two perturbations in different cell type contexts [11]. Including multiple cell types in the training of GEARS could result in compromised predictions.

In conclusion, we offer several recommendations for the practical application of CellCap. Firstly, it is advisable to determine the desired resolution of variations. If the focus is on exploring heterogeneous responses at the nuanced level of cell state, it is recommended to narrow down to a single cell type during the training of CellCap. Conversely, if the objective is to investigate more coarse-grained response heterogeneity at the cell type level, all cell types can be included. Secondly, it is crucial to ensure that the control group of cells encompasses all potential variations in the input cell state. This necessitates a well-designed experiment where control cells and perturbed cells originate from the same population, ensuring that the distribution of control cell states aligns with the distribution of pre-perturbation cell states for the perturbed cells. Lastly, it is important to acknowledge that certain hyperparameter choices, particularly the weight assigned to automatic relevance determination, can impact the interpretation of results. While variations in hyperparameter values may yield slightly different outputs, they also present an opportunity for gaining fresh perspectives on understanding the landscape of perturbation responses. We encourage users to train CellCap with diverse setups and assess the model outputs using domain knowledge and expertise.

## 4 Methods

### 4.1 The CellCap model

CellCap is a probabilistic generative model for single-cell transcriptomics count data resulting from perturbation experiments, where groups of cells are subjected to different perturbation conditions. Perturbation responses are parameterized by linear combinations of learned “response programs”, which can be activated by one or more perturbations. An attention mechanism [17] is used to parameterize a correspondence between the “basal state” of a cell and the perturbation response amplitude, allowing the response to be heterogeneous at the single-cell level.

In designing CellCap, we draw inspiration from existing approaches that model single-cell perturbation data by leveraging latent space arithmetics, in particular the CPA model [12], and reconsider some of their design choices under the lens of model identifiability and interpretability. We pay much attention to where the nonlinearities appear in the model. In the CPA model, a linear latent space algebra gives rise to a latent representation of a cell which is put through a nonlinear decoder. In CellCap, we instead move the nonlinearity from the decoder to the latent space algebra, where an interpretable attention mechanism models the correspondence between cell state and perturbation response. The linear decoder allows us to interpret perturbation responses as the sum of linear “gene expression response programs” – programs which can be unique to one perturbation or shared across several perturbations – and which contribute to the measured response in a cell-state-dependent manner.

#### 4.1.1 The CellCap generative process

A formal definition of the CellCap data generative process is given below, along with a glossary of random variables, intermediate quantities, and the implied meaning of subscript indices of various tensor quantities:

*n* cell *∈* 1 … *N g* gene *∈* 1 … *G*

*p* perturbation *∈* 1 … *N* ^(pert)^

*q* response program *∈* 1 … *Q*

*c* fixed effect covariates *∈* 1 … *N* ^(cov)^

*k* latent space dimension *∈* 1 … *K*

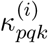*P × Q* learnable *K*-dimensional vectors for each attention head *i*

*P*_*np*_ perturbation “design matrix” indicating perturbation for each cell, values ∈ {0, 1}

*H*_*pq*_ learnable non-negative matrix signifying the usage of response programs per perturbation

*v*_*nq*_ value vectors which sum the relevant response programs for each cell

*D*_*nc*_ *N × N* ^(cov)^ fixed effect “design matrix” indicating fixed effects for each cell

*y*_*ck*_ *N* ^(cov)^ *× K* learnable matrix (fixed effect covariates)

*h*_*nq*_ *N × Q* usage matrix of response programs

*w*_*qk*_ *Q × K* learnable matrix of response programs in the latent space ∈ (−1, 1) interval

*x*_*ng*_ *N × G* observed count data ∈ {0, 1, 2, …}

*ℓ*_*n*_ library size of cell *n*

*α*_*q*_ the Laplace ARD prior scale factors ∈ (0, 1) interval

Φ_*g*_ negative binomial overdispersion of gene *g*

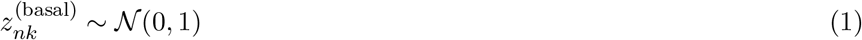

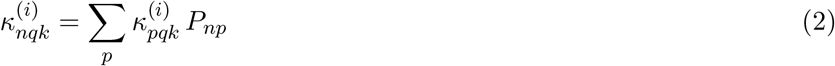

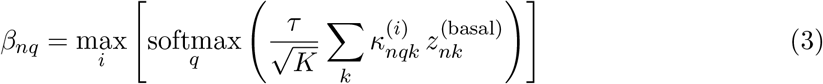

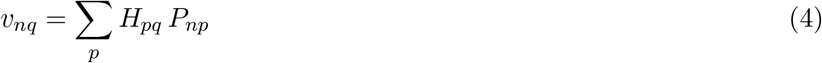

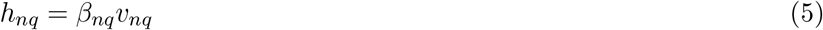

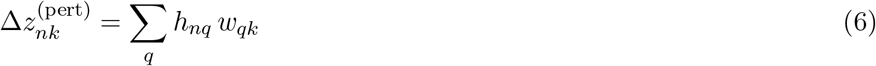

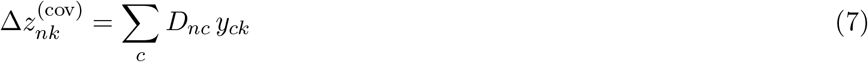

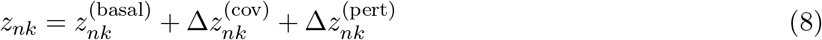

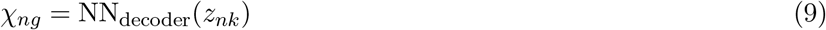

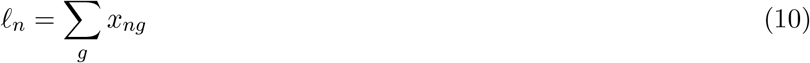

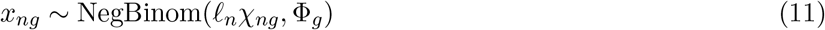

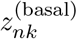 is a *K*-dimensional latent variable representing the “basal” (i.e. unperturbed) state of each cell, with a standard normal prior (Eqn. 1). The final latent representation of a cell, *z*_*nk*_, is recomposed as the sum of 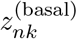 together with two “correction” terms: one for the perturbation itself, 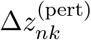, and another for for fixed linear covariates,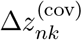. The simpler of these two terms, 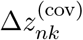, is the matrix product of the covariate design matrix *D*_*nc*_ and the learnable linear fixed effects matrix *y*_*ck*_. Linear fixed effects denoted in the design matrix *D*_*nc*_ can be used to effectively “regress out” technical variation or other variation that is not the subject of the study, such as batch or donor (Eqn. 7).

The perturbation effect, 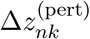, is modeled as a sum over independent response programs *w*_*qk*_ according to their usage by each cell, *h*_*nq*_ (Eqn. 6). *h*_*nq*_ is obtained as the product of a multi-head, scaled dot-product attention mechanism [17], whereby the effects of a given perturbation are made to depend upon the basal state of the cell. This is depicted graphically in Fig. 5. In the language of dot-product attention, the “keys” are *κ*_*nqk*_, the “queries” are 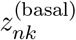, and the “values” are *v*_*nq*_. The keys are derived from the action of a learnable tensor *κ*_*pqk*_ on the perturbation design matrix *P*_*np*_. Here, *κ*_*pqk*_ can be thought of as a dictionary containing *p × q* vectors in the *K*-dimensional latent space for each attention head *i. β*_*nq*_ represents the attention weights that quantify how much cell *n* attends to response program *q* (Eqn. 3). *τ* is a temperature hyperparameter which controls the sharpness of the distribution of attention weight over programs, and by default its value is 4 so that the factor 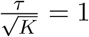 when *K* = 16, the default size of the latent space. The “value” vectors *v*_*nq*_ amount to picking out the response programs for the given cell’s perturbation(s) and summing them (Eqn. 4).

**Figure 5.**
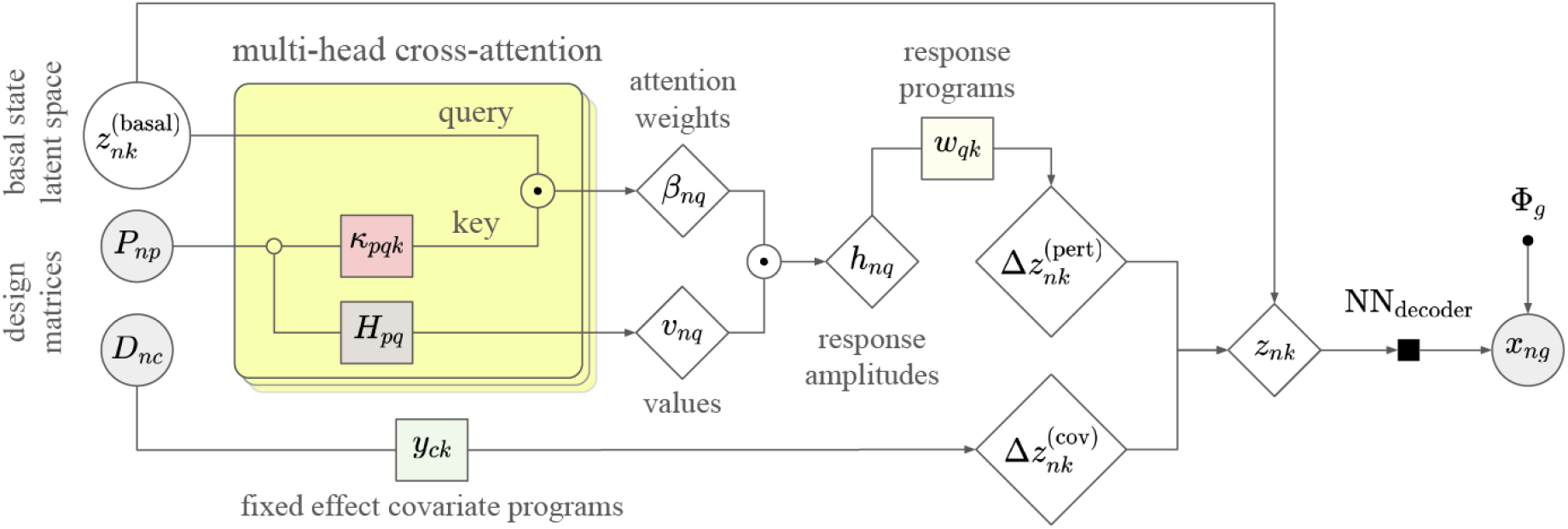
Probabilistic graphical model for the data generative process in CellCap. Observed data are depicted by shaded gray circles (*P*_*np*_, *D*_*nc*_, *x*_*ng*_). Latent variables are in open circles 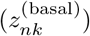. Diamonds are deterministic computations 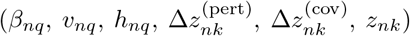. Rectangles of different shadings represent learnable parameters, as do nodes with dots (*κ*_*pqk*_, *H*_*pq*_, *y*_*ck*_, *w*_*qk*_, Φ_*g*_). NN_decoder_ is a single-layer decoder neural network.

Much like the motivation for using multi-head attention (MHA) in natural language processing to capture different semantic views of the words in a given context, we interpret MHA here as a mechanism to capture the one-to-many correspondences between one basal state and different transcriptional response programs. The dot-product attention is computed separately for each head, and for each *n* and *q*, the highest attention score is taken across all heads (Eqn. 3). We deviate from the standard MHA implementation in using the max pooling operation to reduce the effect of multiple heads whereas the original implementation uses a linear combination.

The learned matrix *w*_*qk*_ can then be directly interpreted as latent-space response programs, which can be decoded to gene response programs via the linear decoder, i.e. NN_decoder_(*w*_*qk*_). As discussed by Svensson *et al*., the single-layer decoder lends itself to model interpretation at the expense of a small increase in the reconstruction error [16].

Finally, the complete-information latent space vector *z*_*nk*_, the sum of basal state and perturbation response, is put through the linear decoder NN_decoder_ to obtain *χ*_*ng*_, the normalized gene expression per cell (Eqn. 9). The count data itself, *x*_*ng*_, is sampled from a negative binomial distribution with mean *χ*_*ng*_ times the library size *ℓ*_*n*_ and with a learnable gene-specific overdispersion Φ_*g*_ (Eqn. 11) [14, 15]. As shown in Ref. [25], the zero observations in scRNA-seq data can be effectively accounted for by a negative binomial with appropriately tuned overdispersion without needing to resort to zero inflation to add artificial dropout.

#### 4.1.2 The CellCap variational posterior

We fit CellCap using variational inference [26]. The posterior of CellCap’s only latent variable, 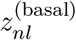, is approximated as:

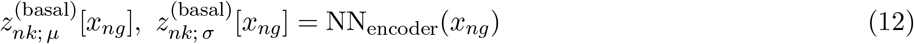

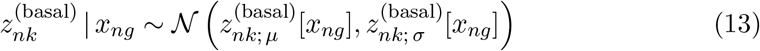

Here, we use a neural network to propose to parameterize the posterior distribution 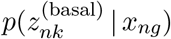, as in Kingma and Welling [26]. Surmising the complexity of the task of inferring the basal state from expression data, we use a deep neural network to this end. We note that this particular inference task is not constrained to be interpretable, justifying our use of a deep neural network.

#### 4.1.3 Posterior regularization via adversarial classification

As at stands, the current modeldoes not include a mechanism to ensure that the learned posterior distribution 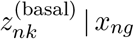 is indeed devoid of perturbation effects. Satisfying this condition, either rigorously or in approximation, is crucial for the correct behavior of CellCap and the interpretation of its results: if perturbation effects, either entirely or in part, are subsumed by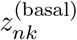, the residual variation that is subject to explicit modeling via 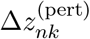, would become non-existent or diminished.

As in the CPA model [12], we use an adversarial classification task to strip perturbation information out of the 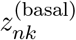 latent space. We use a neural network, NN_classifier_, to try to predict whether a given perturbation *p* was applied in cell *n, P*_*np*_ ∈ {0, 1}. This matrix *P*_*np*_ is the known design matrix given to CellCap as an input. We do the same for each of the covariates supplied in the covariate design matrix *D*_*nc*_. We note that the inclusion of an adversarial loss can be formally related to the theory of posterior regularization by interpreting the adversarial loss term as a KL divergence, see Ref. [27, 28].

Unlike the CPA approach which uses a two-step minimax adversarial gradient update procedure, we apply a gradient reversal layer [29] as the first layer of our adversarial classifier, and we train the whole model with a single gradient update step. Due to the gradient reversal layer, the gradient updates that get applied to the adversarial classifier NN_classifier_ encourage the classifier to perform better classification, while at the same time, the gradients that get applied to NN_encoder_ work against the classifier. This process effectively strips perturbation condition information out of the basal latent space.

#### 4.1.4 Posterior regularization via automatic relevance determination

We additionally include a sparsity-inducing loss term as a form of automatic relevance determination (ARD) [18], inspired by the sparse Bayesian learning literature:

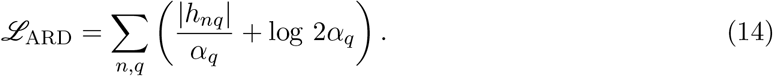

This loss can be construed as the negative log likelihood of *h*_*nq*_ under a Laplace prior with zero mean and scale *α*_*q*_. Intuitively, *α*_*q*_ determines the magnitude of response to program *q*. Performing maximum likelihood estimation over *α*_*q*_ induces sparsity in a data-driven fashion: all things being equal, the data likelihood under the model would be higher if it were to consolidate responses into a few programs *q* with nonzero *h*_*nq*_ by sending certain entries of *α*_*q*_ → 0.

We note that sparsity in the usage of response programs can be induced by placing an ARD prior on either *h*_*nq*_ or *w*_*qk*_, and that to some extent this choice is arbitrary. So long as one of these variables is bounded, a shrinkage prior on the other performs model selection. In our case, we put the shrinkage prior on *h*_*nq*_ and bound *w*_*qk*_ ∈ (−1, 1) interval. This way ℒ_ARD_ ∝ *𝒪* (*N*) so that the hyperparameter *γ* (below) is independent of the size of the dataset, since all pieces of the loss function scale as 𝒪 (*N*).

#### 4.1.5 The full loss function

In CellCap, the loss function is a sum of the variational evidence lower bound (ELBO) and the aforementioned adversarial loss, which acts as a posterior regularization to strip perturbation information out of the basal latent space. We include tunable hyperparameters *α, β*, and *γ* on various terms in the loss function (see below):

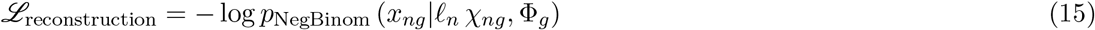

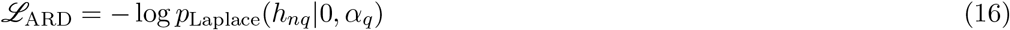

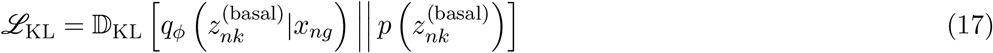

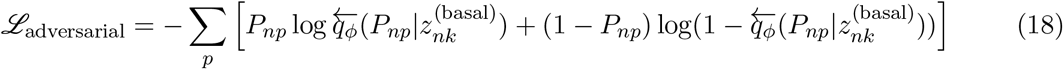

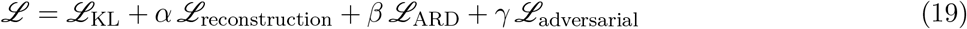

Here, 𝔻_KL_[·||·] is the Kullback-Leibler divergence, and ℒ_adversarial_ is a binary cross entropy loss summed over all perturbations. 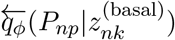 is the posterior binary probability distribution for *P*_*np*_ ∈ {0, 1} (i.e. whether or not perturbation *p* was applied to cell *n*), conditioned on the learned basal cell state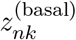. Note that the left arrow indicates the application of a gradient reversal layer as the first layer of the classifier network. The *ϕ* in *q*_*ϕ*_(·) denotes the bundle of learnable parameters {NN_encoder_, NN_classifier_}.

#### 4.1.6 Key model hyperparameters

Each of the four main terms in the full loss function promotes a distinct learning objective. In order to provide users with the flexibility to prioritize different learning objectives as needed, we equip each term with a tunable coefficient as model hyperparameters. These include *α* ≥ 0 coefficient to control *ℒ*_reconstruction_, *β* ≥ 0 coefficient to control *ℒ*_ARD_, and *γ* ≥ 0 coefficient to control *ℒ*_adversarial_. (We note that one of the four loss terms can have its coefficient set to one without loss of generality, so here we have arbitrarily fixed the coefficient of *ℒ*_KL_ to one.) While the choice of proper hyperparameters can vary depending on the dataset, we provide rough guidelines for their tuning based on our interpretation of the role of different terms in the total loss function. The default values for these hyperparameters in CellCap are *α* = 2.0, *β* = 0.2, and *γ* = 1.0. However, we strongly advise users to explore the effects of varying these choices on their results.

Promoting *ℒ*_adversarial_ by increasing *γ* will encourage learning a well-mixed basal state representation. Supplementary Fig. S19 shows an example illustrating how small choices of *γ* can impair the model’s ability to reach a decent fit, as illustrated by the lack of mixing in the left (*γ* = 0) and middle (*γ* = 0.1) columns. Insufficient mixing in the basal state implies disrupting the required compartmentalization of information needed for explicit modeling of Δ**z**^(pert)^. As expected, we notice that the sought after response programs and their correct usage patterns only emerge for *γ* = 1 (right panel). We recommend users evaluate the receiver operating characteristic (ROC) curve of the adversarial classifier to confirm if the model has reached a well-mixed solution for the basal state. An acceptable fit of the basal state encoder should return adversarial classifier ROC curves close to the diagonal line, such that cells from different perturbations are indistinguishable.

Promoting *ℒ*_reconstruction_ decreases the reconstruction error in decoding the recomposed latent representation 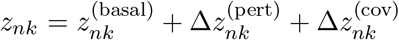 back to the original gene expression profile *x*_*ng*_. Given the influence of 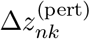 in shaping the recomposed latent representation *z*_*nk*_, promoting the reconstruction loss term additionally drives learning a better and more nuanced description of perturbations, often by learning additional response programs. We control the sparsity of the response programs via the *ℒ*_ARD_ term with strength controlled by the *β* coefficient. Therefore, *α* and *β* play conflicting roles. In practice, we have noticed that choosing the appropriate *α* and *β* varies significantly from one dataset to another. For improved perturbation prediction, users can increase *α* while decreasing *β*. However, doing so may render *w*_*qk*_ unnecessarily complicated and difficult to interpret. Alternatively, increasing *β* will result in sparser transcriptional programs, as demonstrated in Supplementary Fig. S20. Naturally, choosing very large values of *β* poses the risk of attaining an over-simplified fit.

#### 4.1.7 Key implementation details

The CellCap model is implemented in Python on top of the `scvi-tools` framework for probabilistic modeling of single-cell data [30]. The `scvi-tools` framework streamlines the implementation of models following the setup of Kingma and Welling’s variational autoencoders for Bayesian inference [26] and provides much of required logic for single-cell data loading and model training, leveraging the PyTorch Lightning library. The reconstruction loss is evaluated by using single Monte Carlo posterior samples of *h*_*nq*_ and 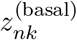 to compute *χ*_*ng*_ in the generative model. The adversarial classifier is implemented as a dense neural network with 2 hidden layers and 1 output layer. Both hidden layers have 128 neurons, and the last layer predicts a probabilistic perturbation assignment matrix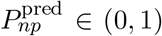. A gradient reversal layer is inserted before the first hidden layer, for the purpose of adversarial training.

In practice, we found it quite beneficial for rapid model convergence to carefully initialize the tensor 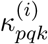 to a set of representative points from 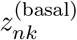. We choose these representative points using Louvain community detection to identify *Q* random “key” cells from the control group, using their initial basal states as 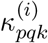 for each perturbation and attention head. This ensures that the key vectors start out in a configuration inside the distributional support of the basal state. Poor initialization of 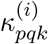 results in vanishing gradients due to the softmax operation and slow convergence.

For datasets that have extreme class imbalance, we implemented a weighted random sampler to balance the ratio of control and perturbed cells in each mini-batch during training. The sampling weight for each condition was calculated as *N/*(*N*_classes_ *× N*_*c*_), where *N* is the total number of samples, *N*_classes_ is the number of classes, and *N*_*c*_ is the number of samples in class *c*. This calculation is implemented in `sklearn.utils.class weight.compute sample weight`. Training of CellCap is equipped with NVIDIA Tesla T4 GPU. We use the AdamW optimizer to update all trainable weights in the CellCap model. The initial learning rate is set to 10^−3^ followed by reductions by a factor of 0.6 after each plateau, as implemented in PyTorch’s `ReduceLROnPlateau` learning rate scheduler. We implemented early stopping, which is triggered if the loss does not reduce for more than 50 epochs. Otherwise, training stops after reaching the maximum number of epochs, which is 1000 in all cases in this study.

The number of cells and the number of perturbations vary across single-cell perturbation datasets, which can influence the the total runtime of CellCap. For the real perturbation datasets used in this study, the total runtime ranges from 1 - 2 hours on a single Nvidia Tesla T4 GPU.

## Data Availability

Code to simulate all three scenarios and their ready-to-use data can be found at https://github.com/broadinstitute/CellCap. Raw data for pathogen-exposure human monocytes is deposited at the European Genome-Phenome Archive (EGAS00001005376). A processed version is hosted at https://eqtlgen.org/sc/datasets/1m-scbloodnl.html. Raw data from Norman *et al*. is deposited at Gene Expression Omnibus (GEO), under accession number GSE133344. A processed version of the Norman *et al*. data can also be found at http://projects.sanderlab.org/scperturb[31].

## Code Availability

CellCap code and quick-start tutorials are available at https://github.com/broadinstitute/CellCap. To reproduce major results in this study, please follow the notebooks provided in the CellCap GitHub repository.

## Acknowledgements

This work was supported by the Broad Institute’s Stanley Center for Psychiatric Research, Data Sciences Platform, and the National Institute of Mental Health (grants U01MH115727 and UM1MH 130966). We thank Jim Nemesh and Noah Pettinari for participating in discussions and providing insightful feedback throughout the project.

