## Supplementary materials for "Modeling interpretable correspondence between cell state and perturbation response with CellCap"

### S Supplemental Material

#### S.1 Generating simulated perturbation scRNA-seq datasets

Here, we provide additional details regarding the simulated perturbation scRNA-seq datasets discussed in the main text. In all cases, we generated 4 cell types (A, B, C, and D) using PROSSTT [19] along a continuous differentiation process. Cells in each cell type are positioned along a continuum of states according to their pseudotime $t \in [0, 1]$ parameter. We used the expression profile of cell type A as the basal state (also to serve as the unperturbed control), while treating expression profiles of cell type B, C and D as 3 perturbation programs. For consistency across all simulation scenarios, we chose the expression profiles of cell type B and C as perturbation-specific programs and expression profile of cell type D as the shared response program.

For concreteness, we consider the expression of cell type B and treat it as a fictitious response program. Other simulated scenarios follow similar logic. We generated perturbed gene expression counts from the following stochastic process.

$$x_{ng}^{(\mathrm{pert})}(t) = x_{ng}^{\mathrm{A}}(t) + \Delta x_{ng}^{(\mathrm{pert})}(t)$$
$$\Delta x_{ng}^{(\mathrm{pert})}(t) \sim \mathrm{Poisson}\left(\lambda_{ng} = C\,t\,\frac{x_{ng}^{\mathrm{B}}}{\sum_g x_{ng}^{\mathrm{B}}} + C\,(1-t)\,\frac{\sum_n x_{ng}^{\mathrm{A}}}{\sum_{ng} x_{ng}^{\mathrm{A}}}\right),$$

where $C$ is the average library size associated with the perturbation effect, which we set to the average library size of cells in group A (unperturbed). The second Poisson rate term is to counteract the extra library size proportional to $t$ and is simply adjusting the expression with a non-informative sample from the basal state pseudo-bulk with library size $1 - t$. The final library size is $\bar{\ell}_A + C$ on average, where $\bar{\ell}_A$ is the average library size of cells in group A (unperturbed).

Besides the three simulation scenarios presented in the main text, we included three more simulation scenarios for extended scrutiny. In the fourth scenario, two perturbations induce two unique response programs and one shared response program (Supplementary Fig. S21a). The optimal solution identified by CellCap is decomposing perturbation effects into two distinct programs while ignoring the shared set of perturbed genes. We reasoned that indeed CellCap's operation is justified given the sparsity regularization and the fact that the data provides no support for the shared program being sampled independently of the two unique programs.

The results from the other two scenarios are presented in Supplementary Fig. S21a. In the fifth scenario, to break the tie present in the fourth scenario, we introduced a third perturbation inducing only the shared response of the first two perturbation. In the sixth scenario, we simply generated more cells from perturbation conditions to provide CellCap with more data to learn the programs and their shared versus unique usage. Training CellCap under these additional scenarios shows that: (1) CellCap will not distinguish a shared response program as an independent program if there is no evidence to establish the shared program as an independent program (as in the fourth scenario), and (2) the shared program will become distinguishable from the unique programs provided enough cells support all three transcriptional programs as independent programs (Supplementary Fig. S21b).

#### S.2 Average perturbation response

There are two quantities in CellCap which can be used to convey the average response of cells in a given perturbation condition, and they have slightly different meanings. $H_{pq}$ represents the maximum possible usage of each response program by each perturbation (since in Eqn. 5, $\beta_{nq} \in (0, 1)$). $H_{pq}$ does not explicitly take the basal state into account. $h_{nq}$, on the other hand, quantifies the per-cell usage of each response program, conditioned on the applied perturbation and the basal cell state. For that reason, $h_{nq}$ gives a more accurate picture of cellular responses. When we compute the "average perturbation response", we compute the average $\frac{1}{N_p}\sum_{n \in \mathcal{G}_p} h_{nq}$ over groups of cells $\mathcal{G}_p$ in each perturbation condition $p$, where there are $N_p$ cells in group $\mathcal{G}_p$.

#### S.3 Identification of basal expression patterns associated with response programs

Identifying the most relevant basal state that elicits a response program is similar to the process of matching a perturbation key $\kappa_{pq}$ with basal state $\mathbf{z}_n^{(\text{basal})}$. Given a response program of interest together with its latent space key $\kappa_{pq}$, we define its "*relevance score*" as the cosine similarity to each basal state $\kappa_{pq} \cdot \mathbf{z}_n^{(\text{basal})}$. We note that given the dot-product attention mechanism, cosine similarity is the optimal similarity metric within the framework of CellCap. Finally, to identify the top $M$ marker genes that best correlate (or anti-correlate) with the program relevance score, we fitted a univariate linear regression using log-normalized gene expression profile $x_{ng}$ as the input and relevance score as the target value. We additionally performed an F-statistic test to report p-values for each gene feature and selected the best $M$ features. We implemented this workflow using `sklearn.feature_selection.SelectKBest` and `sklearn.feature_selection.f_regression` from the scikit-learn package.

#### S.4 Identification of top responding genes associated with each response program

Identification of top responding genes associated with each response program is simply achieved by decoding the learned response dictionary elements (i.e. the rows of $w_{qk}$ matrix) to obtain the $Q \times G$ matrix $w_{qg}$. We note that this operation is made straightforward merely by the virtue of using a linearly-decoded latent space. Had we used a nonlinear decoder (e.g. as in the CPA model), the semantics of latent space translations would become intricately local and cell-state-dependent.

Here, we further demonstrate that the decoded transcriptional programs $w_{qg}$ are closely related to the log-fold-change (LFC) gene expression of perturbed cells compared to basal cells. For simplicity, we consider a simple scenario where $\mathbf{z} = \mathbf{z}^{(\text{basal})} + h\,\mathbf{w}$, corresponding to the contribution of a single transcriptional program with amplitude $h$. According to the CellCap model, we have:

$$\begin{aligned}
\mathbb{E}\left[x_g \,|\, \mathbf{z}\right] &= \ell\,\chi_g \\
&= \ell\,\text{softmax}\left[\sum_k W_{gk}^{(\text{decoder})}\left(z_k^{(\text{basal})} + h\,w_k\right)\right] \\
&= \ell\,\frac{\exp\left(y_g^{(\text{basal})}\right)\,\exp\left(hw_g\right)}{\sum_{g'}\exp\left(y_{g'}^{(\text{basal})}\right)\,\exp\left(hw_{g'}\right)} \\
&\simeq \ell\,\chi_g^{(\text{basal})}\left[1 + hw_g - h\frac{\sum_{g'} w_{g'}\exp\left(y_{g'}^{(\text{basal})}\right)}{\sum_{g'}\exp\left(y_{g'}^{(\text{basal})}\right)} + \mathcal{O}(h^2)\right]
\end{aligned} \tag{20}$$

where we have defined shorthands $y_g^{(\text{basal})} = \sum_k W_{gk}^{(\text{decoder})}\,z_k^{(\text{basal})}$, the decoded program $w_g = \sum_k W_{gk}^{(\text{decoder})}\,w_k$ like before, and $\chi_g^{(\text{basal})} = \exp\left(y_g^{(\text{basal})}\right) / \sum_{g'}\exp\left(y_{g'}^{(\text{basal})}\right)$ as the normalized basal state gene expression. The last line of Eq. (20) is readily obtained by first-order Taylor's expansion by assuming that the perturbation effect is weak compared to the basal gene expression. By definition of LFC, we have:

$$\text{LFC}_g = \frac{\mathbb{E}\left[x_g \,|\, \mathbf{z}^{(\text{basal})} + h\mathbf{w}\right]}{\mathbb{E}\left[x_g \,|\, \mathbf{z}^{(\text{basal})}\right]} \simeq h\,(w_g + \text{const.}) + \mathcal{O}(h^2), \tag{21}$$

where again we have used first-order Taylor's expansion and Eq. (20). As we were hoping to show, the LFC induced by the activation of a program is indeed proportional to the decoded program, save for a constant (the third term in square brackets in Eq. 20). This observation further substantiates the interpretability of the CellCap-derived gene programs $w_{qg}$, and also shows that these decoded programs can be construed as normalized and signed effect sizes with respect to the basal expression, and independent of the basal expression rate of a gene. Given that $w_{qg}$ are normalized and bounded, we can further approximate their distribution as normal, convert decoded gene weights to $z$-scores separately for each program, and then assign $p$-values to each gene using a two-tailed $t$-test to identify which genes have significant effect sizes ("responding") in each program. We define genes with $p$-values $< 0.05$ as responding genes.

#### S.5 Gene set enrichment analysis

We ran GSEA using the web-based tool WebGestalt [32]. For gene ontology annotation, we selected biological processes powered by PANTHER [33]. For pathway annotation, we used the KEGG database [34]. Categories with less than 5 or over 2000 genes were excluded from GSEA. GSEA was performed with 1000 permutations, and results with FDR $\leq 0.1$ were considered to be significant.

#### S.6 Scoring cell cycle

The five stages of the cell cycle are G1/S, S, G2/M, M, and M/G1. Gene sets associated with these five stages have been identified in a previous study [35]. We scored each cell cycle stage by summing log-normalized expressions of all genes in each category [1]. We then standardized the score of each cell cycle stage by subtracting the mean and scaling to unit variance across the entire cell population. Log-normalization was done with Scanpy package,as part of common steps in scRNA-seq data processing[36].

#### S.7 Software packages

Details of all software packages used are included in Supplementary Table S1.

Table S 1: Details of software packages used.

| Package | Version | Reference | URL |
|---|---|---|---|
| CellCap | 0.1.0 | (this work) | `https://github.com/broadinstitute/CellCap` |
| scvi-tools | 0.19.0 | Ref. [30] | `https://scvi-tools.org` |
| PyTorch | 1.11.0 | Ref. [37] | `https://github.com/pytorch/pytorch` |
| Scanpy | 1.9.1 | Ref. [36] | `https://scanpy.readthedocs.io/en/stable` |
| scikit-learn | 1.3.0 | Ref. [38] | `https://scikit-learn.org/stable` |

#### S.8 Supplemental Results

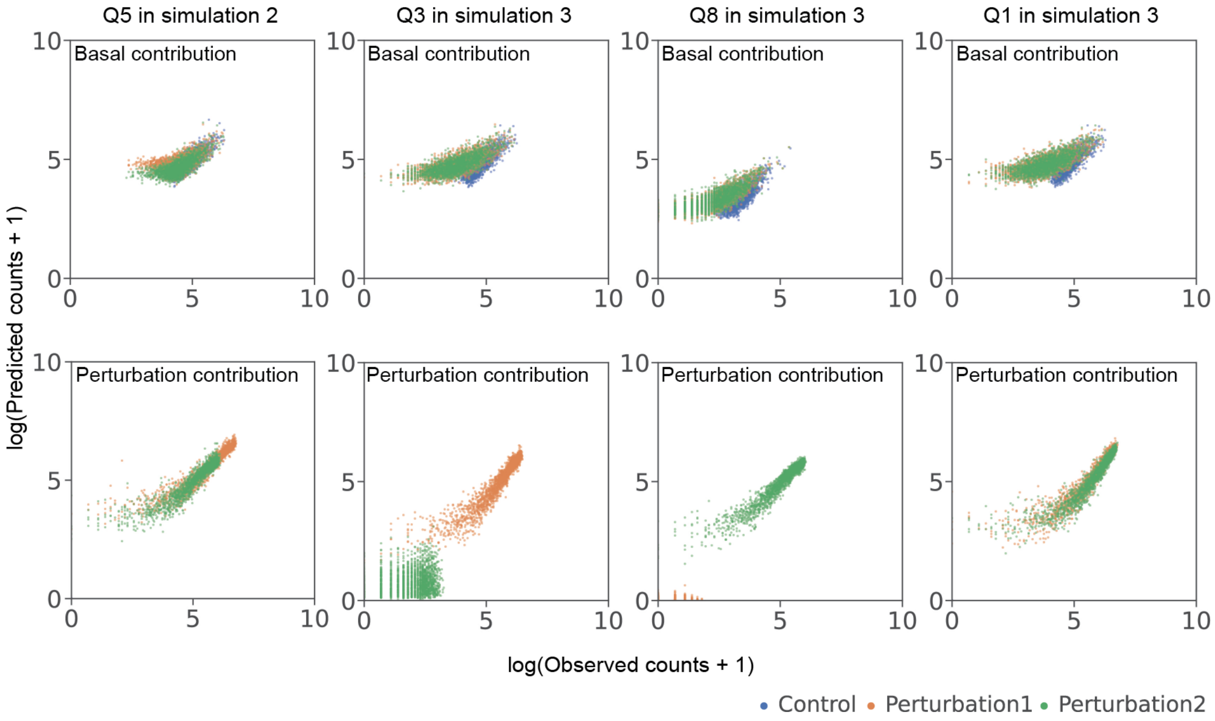


Figure S 1: Total gene expression counts associated with the basal state $\mathbf{z}_n^{(\text{basal})}$ (top) or the perturbation term (bottom) by CellCap in scenario (2) and (3) versus the simulated ground truth. Each dot is a cell, colored by applied perturbation. The perturbation contribution to gene expression (bottom row) is obtained by decoding the final latent space representation $\mathbf{z} = \mathbf{z}^{(\text{basal})} + \Delta\mathbf{z}^{(\text{pert})}$ and then subtracting the contribution of the basal-only latent representation $\mathbf{z}^{(\text{basal})}$.

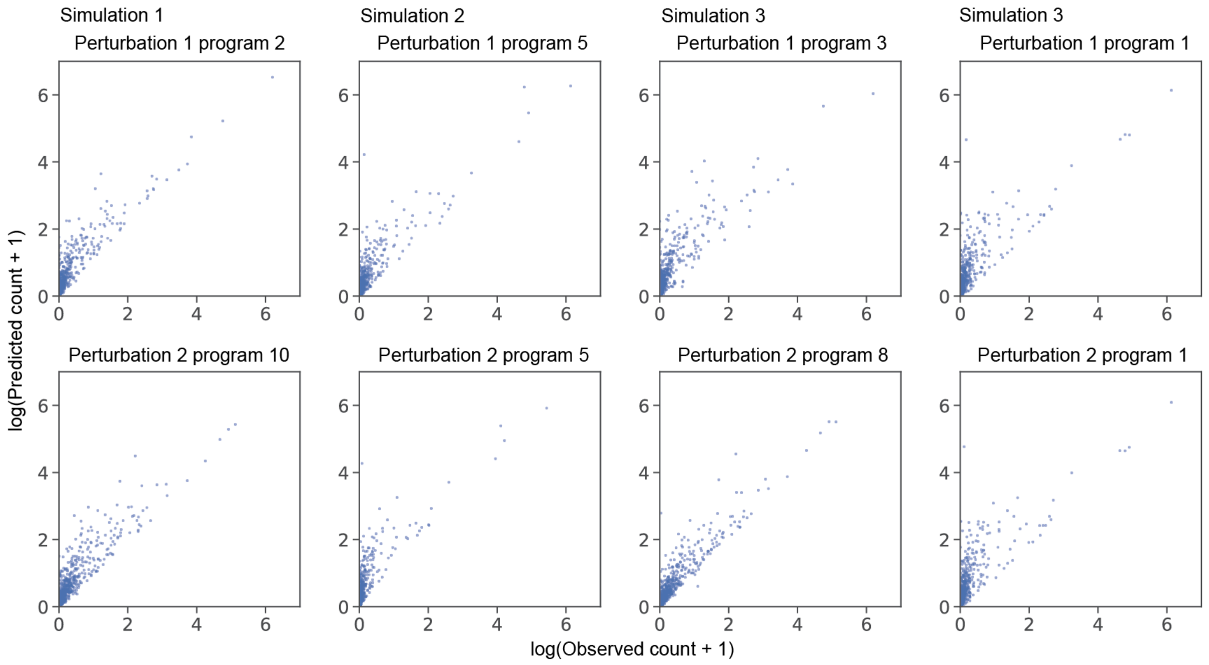


Figure S 2: Gene-wise prediction of CellCap for the simulated scenarios versus the simulated ground truth. Each dot represents the average expression of a gene across all cells in that perturbation condition.

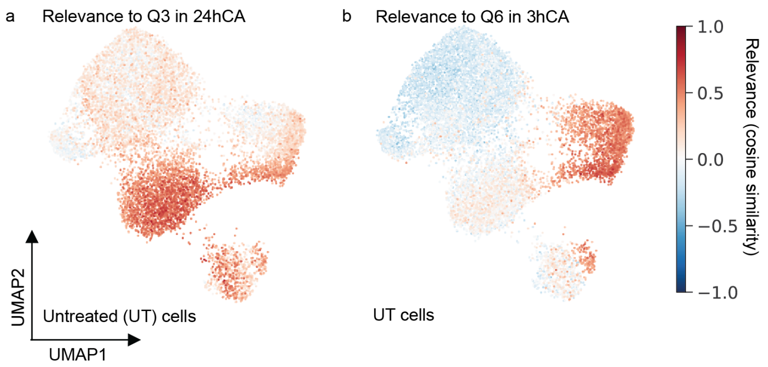


Figure S 3: UMAP plots showing the "relevance score" (Supplementary Section S.3) of untreated cells for the activation of transcriptional programs Q3 (a) and Q6 (b) in the human monocyte data from Fig. 3.

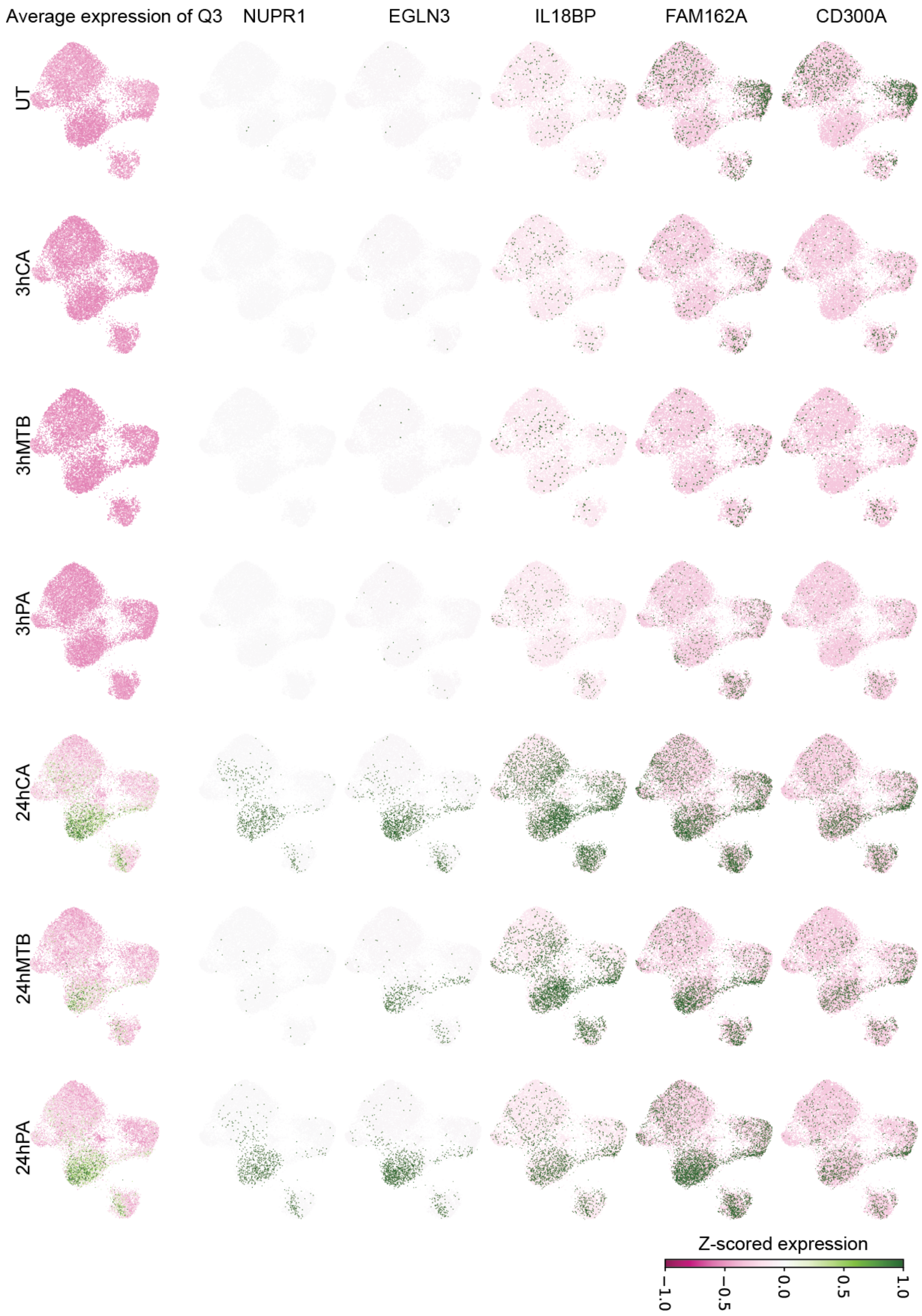


Figure S 4: Basal state UMAPs showing gene expression patterns associated with response program Q3 in the human monocyte data from Fig. 3. Though the basal state UMAP is shown, the gene expression data is $x_{ng}$. The first column shows the average expression pattern of perturbed genes in Q3, and the rest of the columns show the expression pattern of several selected genes from the Q3 program across all conditions.

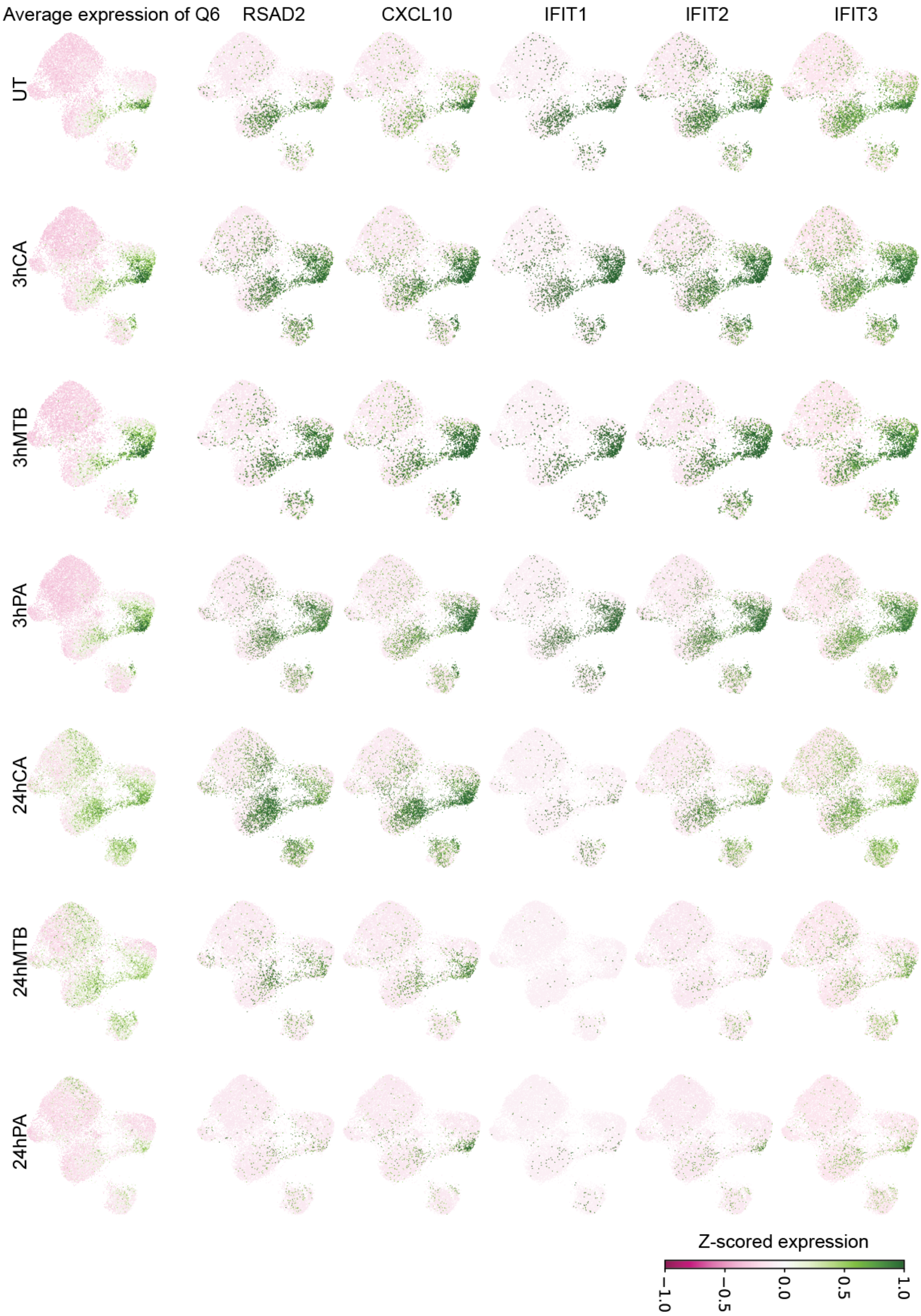


Figure S 5: Basal state UMAPs showing gene expression patterns associated with response program Q6 in the human monocyte data from Fig. 3. Though the basal state UMAP is shown, the gene expression data is $x_{ng}$. The first column shows the average expression pattern of perturbed genes in Q6, and the rest of the columns show the expression pattern of several selected genes from the Q6 program across all conditions.

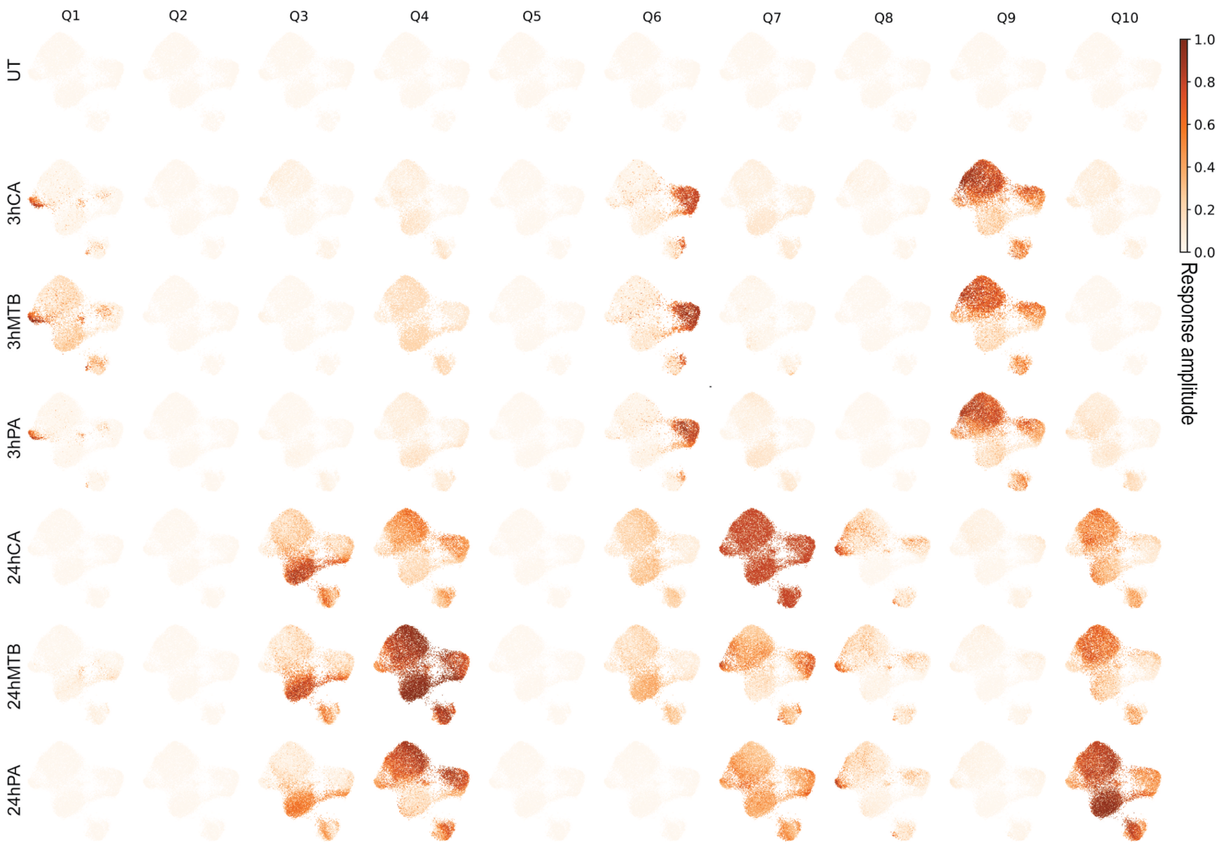


Figure S 6: Basal state UMAPs showing patterns of response amplitudes of 10 transcriptional programs across all 7 conditions in the human monocyte data from Fig. 3.

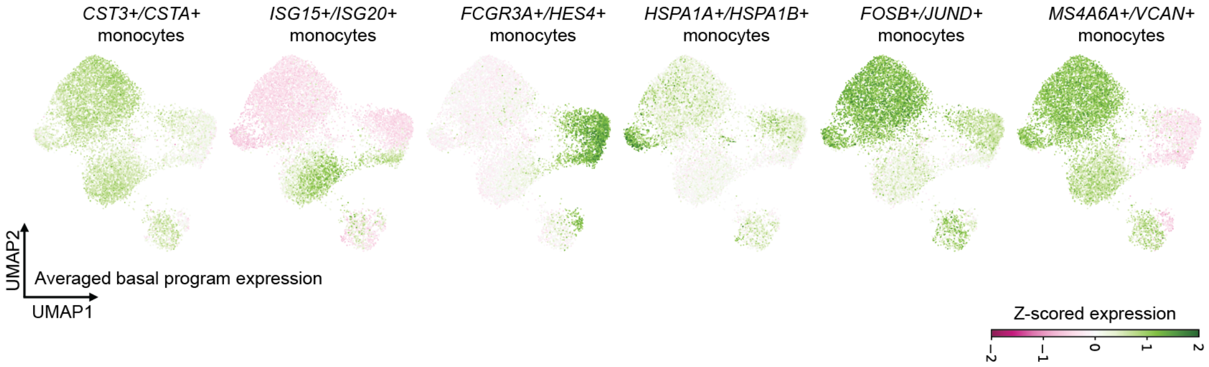


Figure S 7: Soft annotation of human monocytes. UMAPs depict the basal state of control cells from Fig. 3. All response programs shown in 6 were used to identify corresponded basal state programs, and 6 major basal state programs are identified and annotated with two selected genes. Cells are colored by average expressions of basal state programs. Annotations were given by two genes of interest in each basal state program.

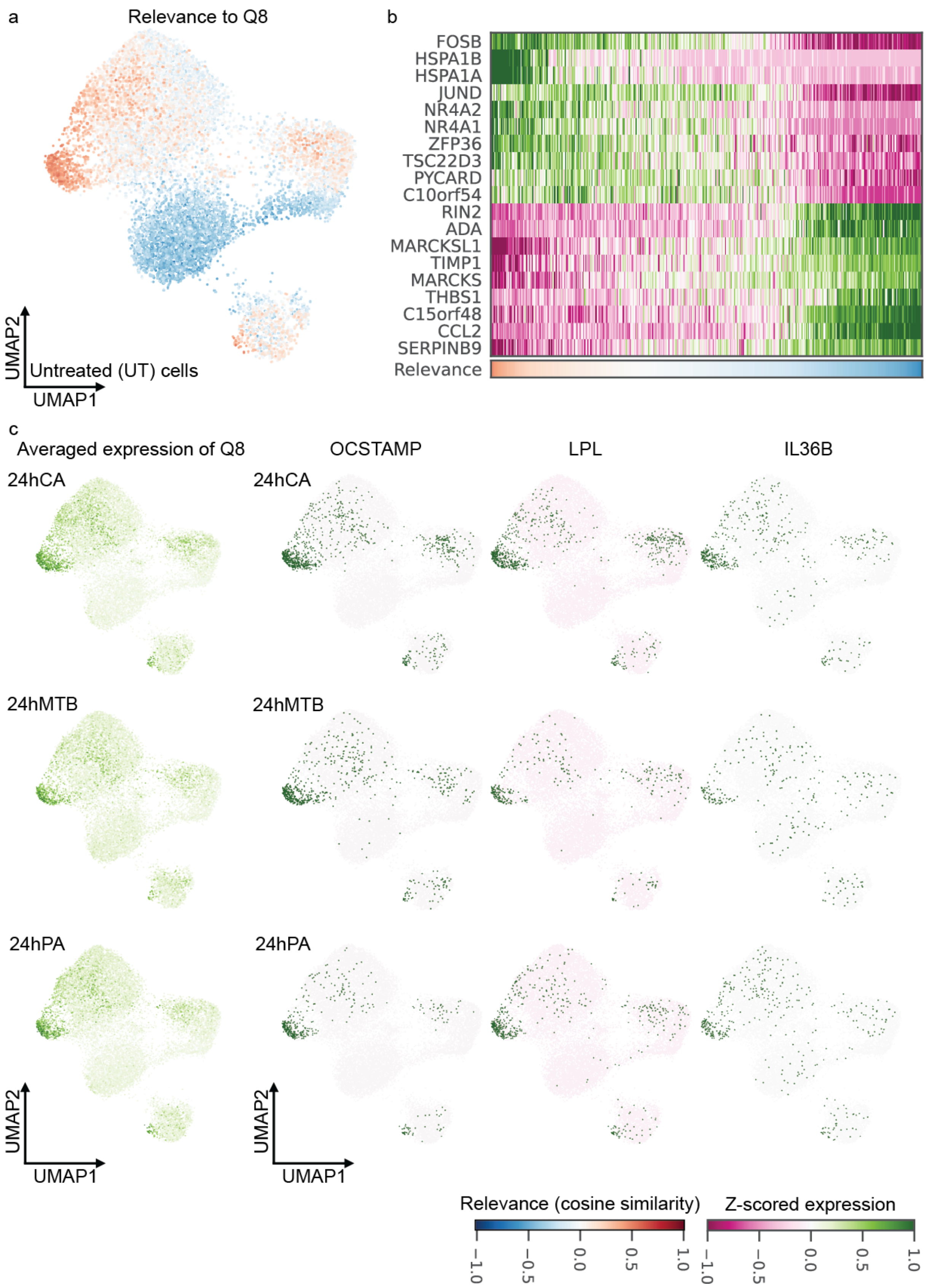


Figure S 8: Basal state UMAPs showing various aspects of response program Q8 in the human monocyte data from Fig. 3. (a) The top left UMAP shows how relevant response program Q8 is to untreated cells, given their basal states. Cells are ordered by highest relevance to lowest in the heatmap on the top right. (b) Top positive and negative basal marker genes are shown. (c) Averaged expression pattern of perturbed genes in Q8 and 3 genes of interest are shown in the bottom UMAP grid.

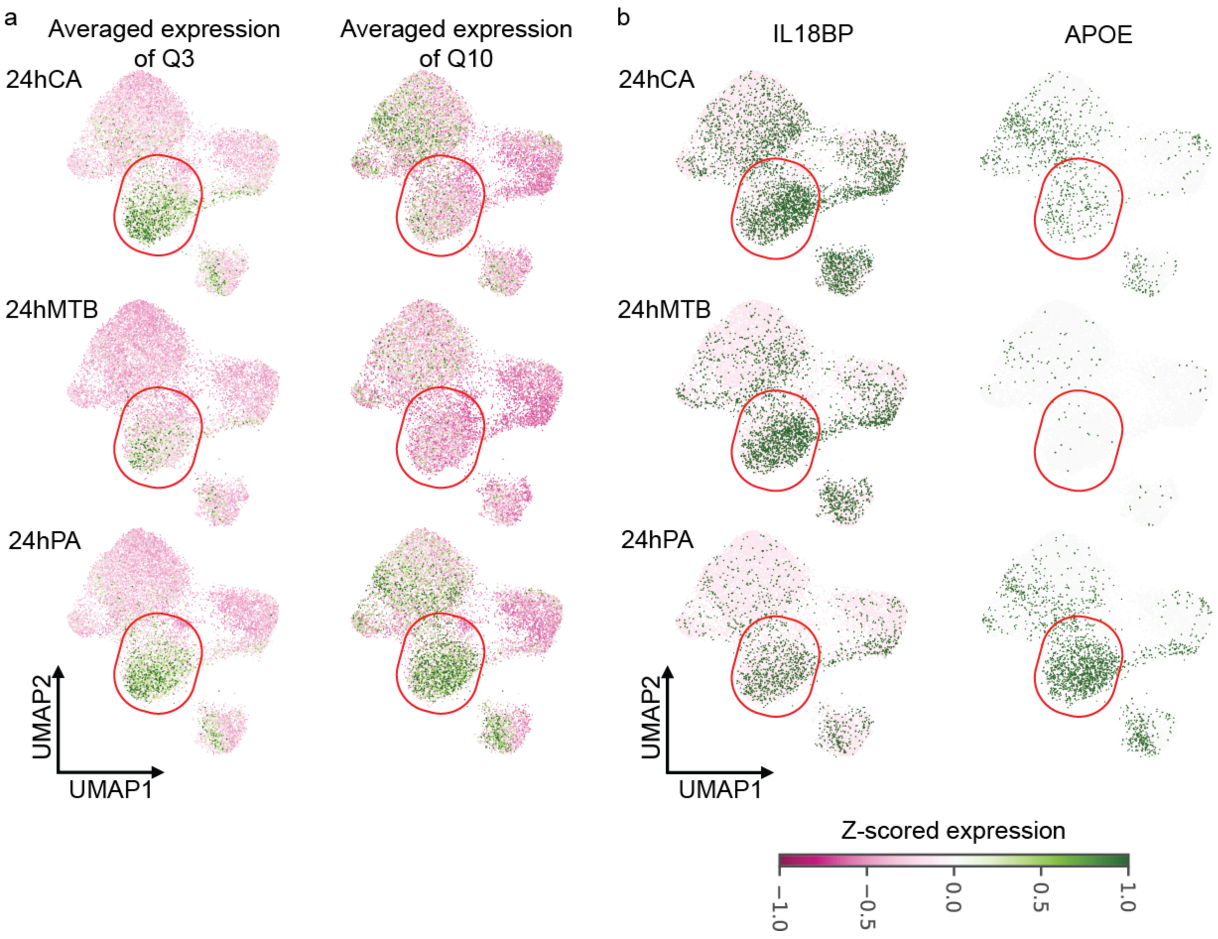


Figure S 9: Comparison of Q3 and Q10 response programs in the human monocyte data from Fig. 3. UMAPs of the basal state are shown. (a) Average expression patterns of responding genes in Q3 and Q10, in conditions of 24hCA, 24hMTB, and 24hPA. (b) Expression patterns of two genes of interest in Q3 and Q10, in 24hCA, 24hMTB, and 24hPA conditions. For (a) and (b), red oval circles highlight the shared and unique expression patterns in these 3 conditions.

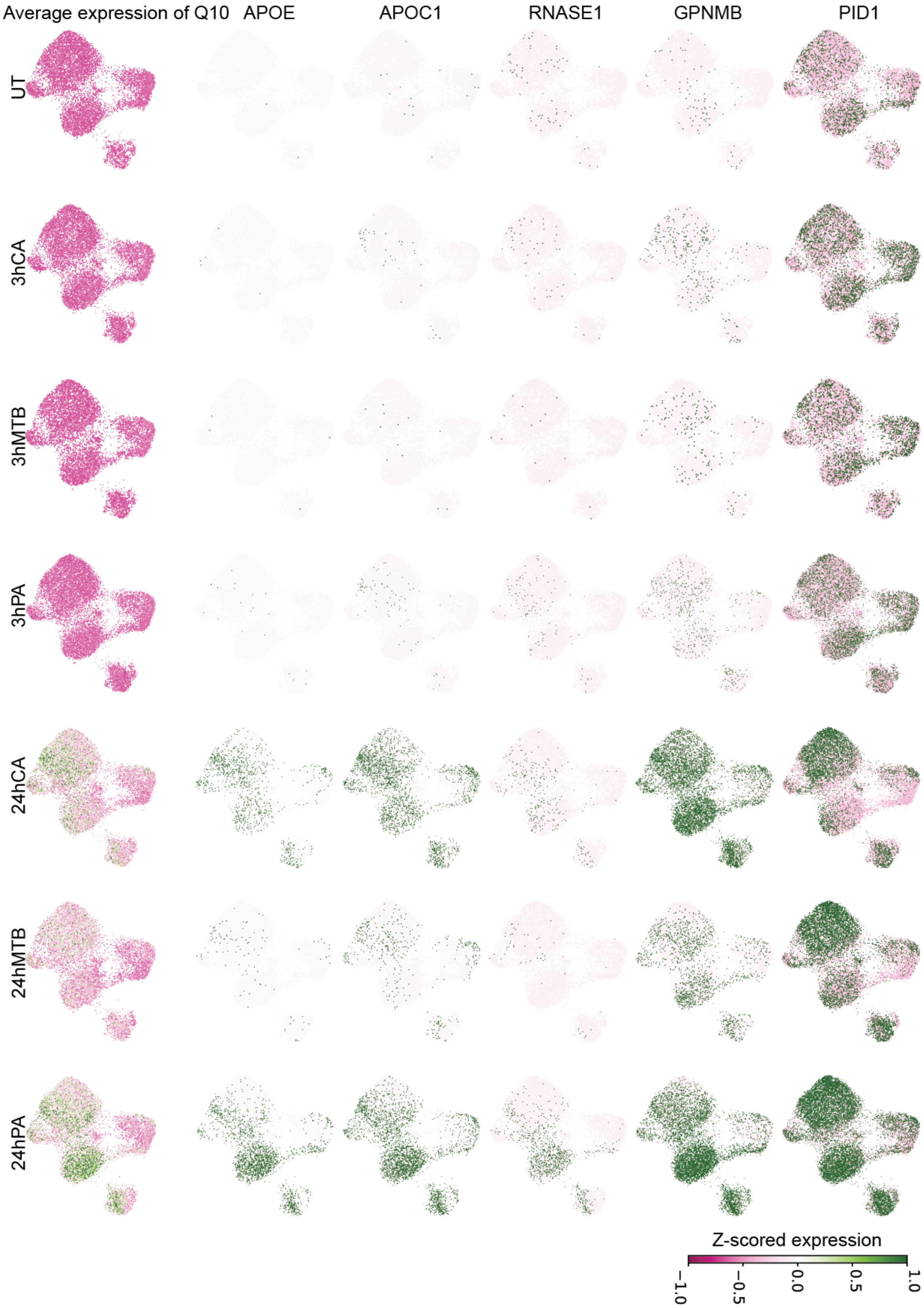


Figure S 10: Basal state UMAPs of gene expression patterns associated with transcriptional response program Q10 in the human monocyte data from Fig. 3. Program Q10 is specific to condition 24hPA. Collective and individual expression patterns of responding genes in Q10 are shown to highlight this specificity. The first column shows the average expression pattern of responding genes in Q10, and the rest of the columns show the expression pattern of several selected genes from the Q10 program across all conditions.

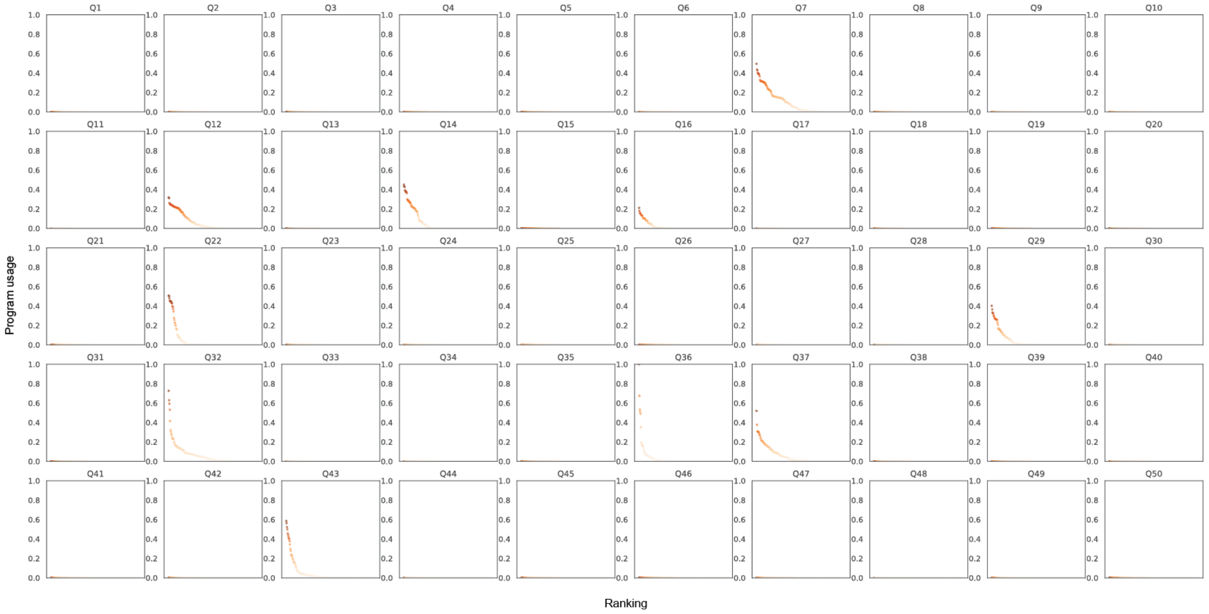


Figure S 11: Rankings of 236 perturbations across 50 response programs in the K562 Perturb-seq data. Program usage of each program is given to each perturbations, and perturbations are sorted from high to low. 10 of 50 programs are shared by 236 perturbation conditions, while the rest of 40 are turned by ARD.

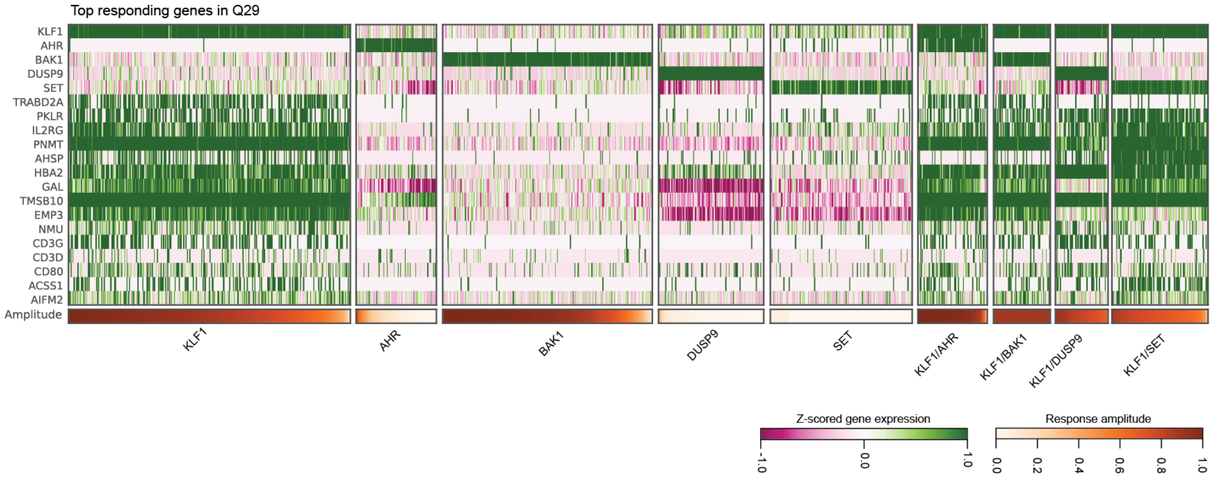


Figure S 12: Top responding genes in transcriptional program Q29 from the K562 Perturb-seq data from Fig. 4. Cells in each condition are ordered from the highest response amplitude to the lowest. Top 5 rows show expressions of CRISPR-targeting genes, and the rest of the rows are genes involved in the Q29 response program.

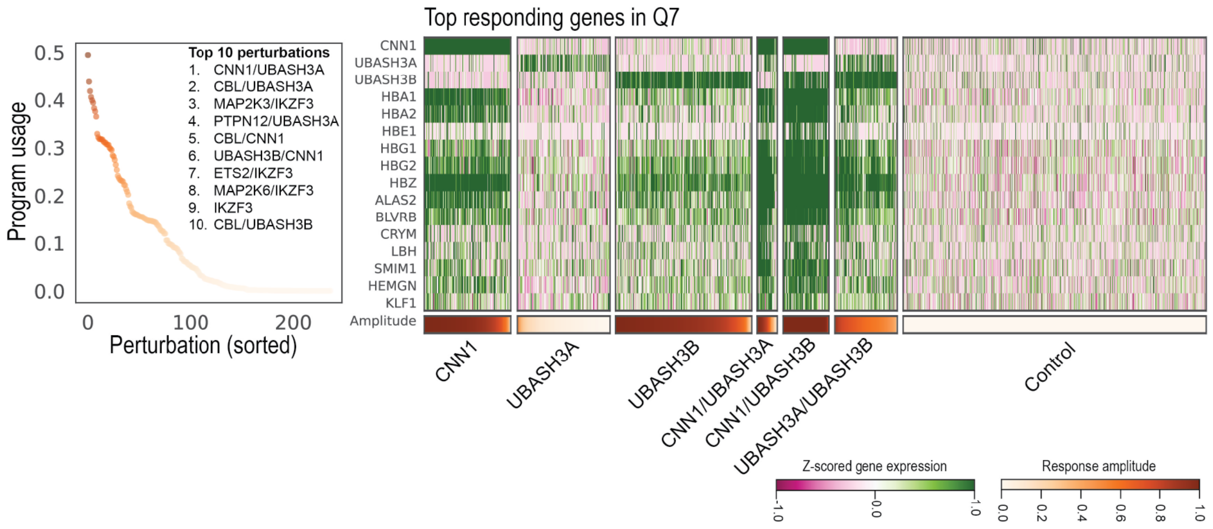


Figure S 13: K562 Perturb-seq data from Fig. 4. Top 10 perturbations associated with Q7 response, and top responding genes in Q7. Cells are ordered from the highest response amplitude to the lowest, and top responding genes are plotted in the heatmap. Top 3 rows show expression of CRISPR-targeting genes, while the rest of the rows show genes involved in the Q7 response program.

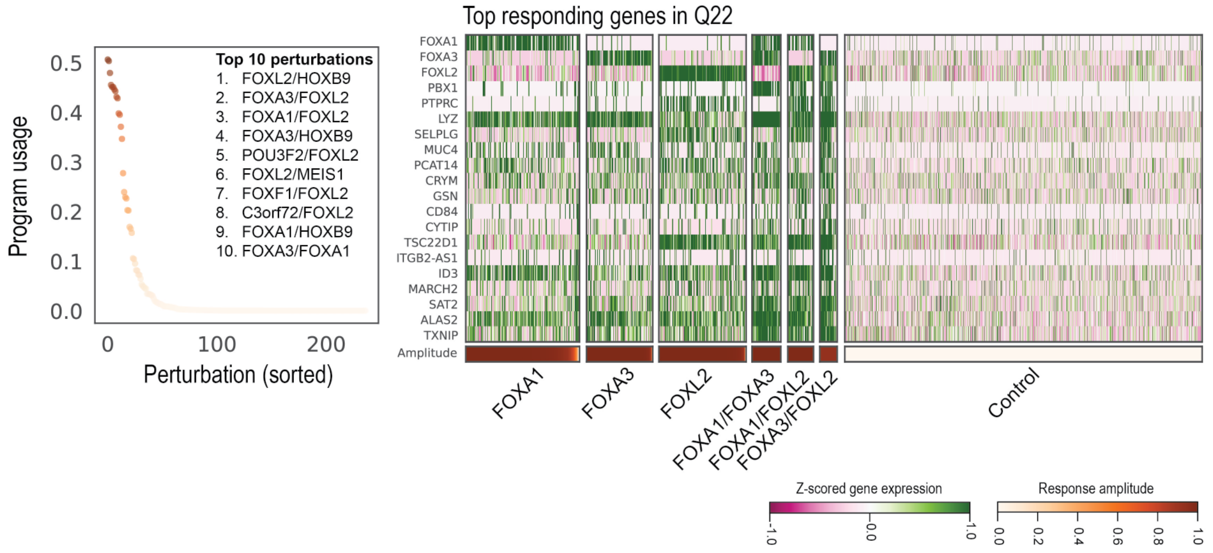


Figure S 14: K562 Perturb-seq data from Fig. 4. Top 10 perturbations associated with Q22 response, and top responding genes in Q22. Cells are ordered from the highest response amplitude to the lowest, and top responding genes in each are plotted in the heatmap. Top 3 rows show expressions of CRISPR-targeting genes, while the rest of the rows show genes involved in the Q22 response program.

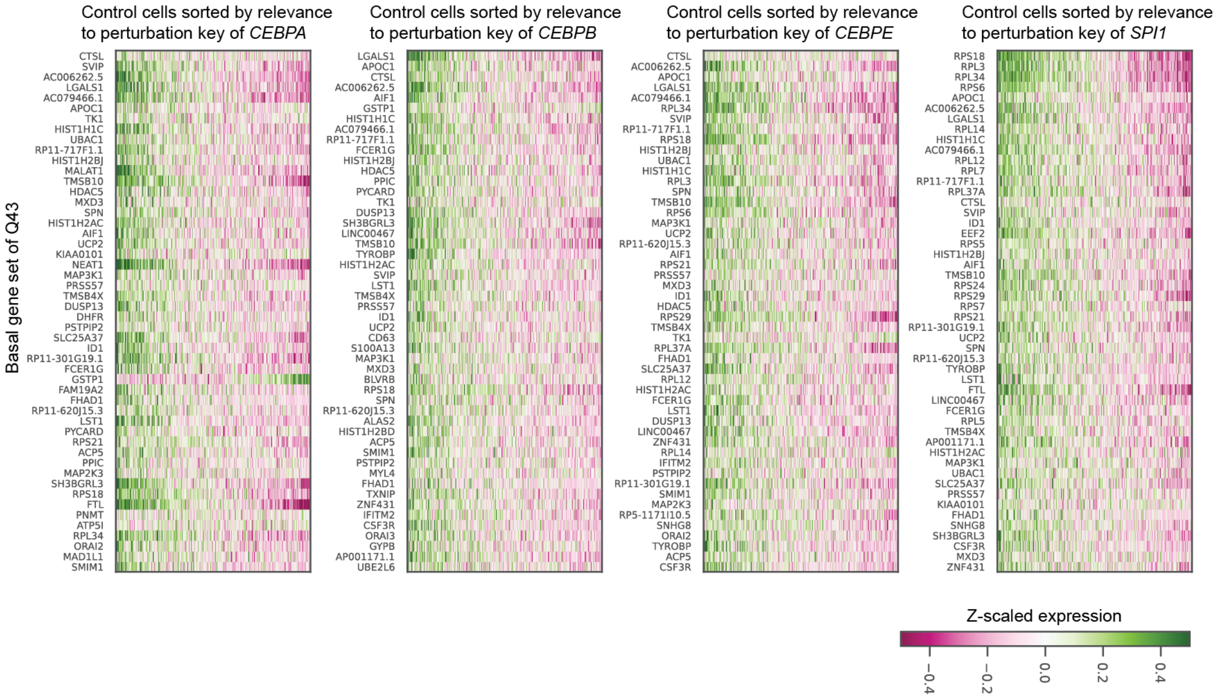


Figure S 15: K562 Perturb-seq data from Fig. 4. Basal state gene expression associated with response program Q43. Cells of control group are ordered by the highest relevance to program Q43 to the lowest. Top positive basal marker genes on both extreme are shown to present the difference of basal expression program. Top 3 rows show expressions of CRISPR-targeting genes instead of being presented as responding genes.

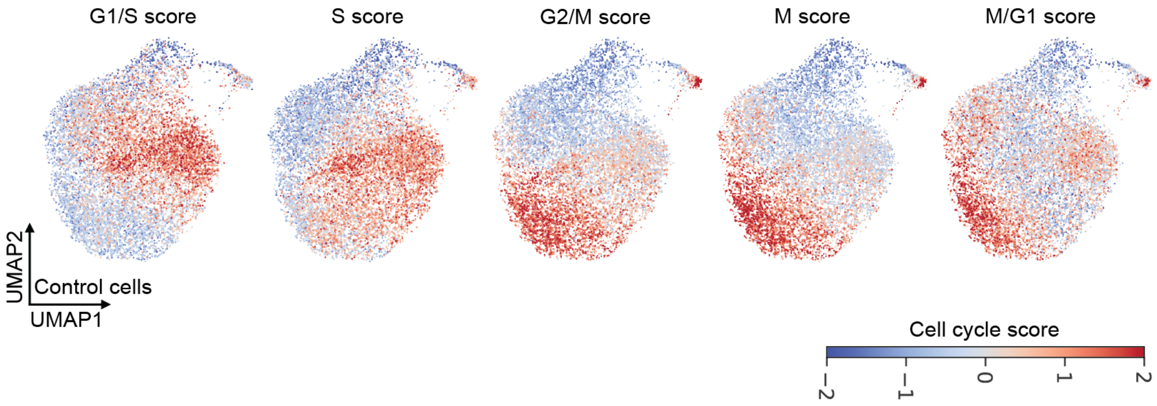


Figure S 16: Scores of 5 cell cycle stages in control cells from the K562 Perturb-seq data from Fig. 4.

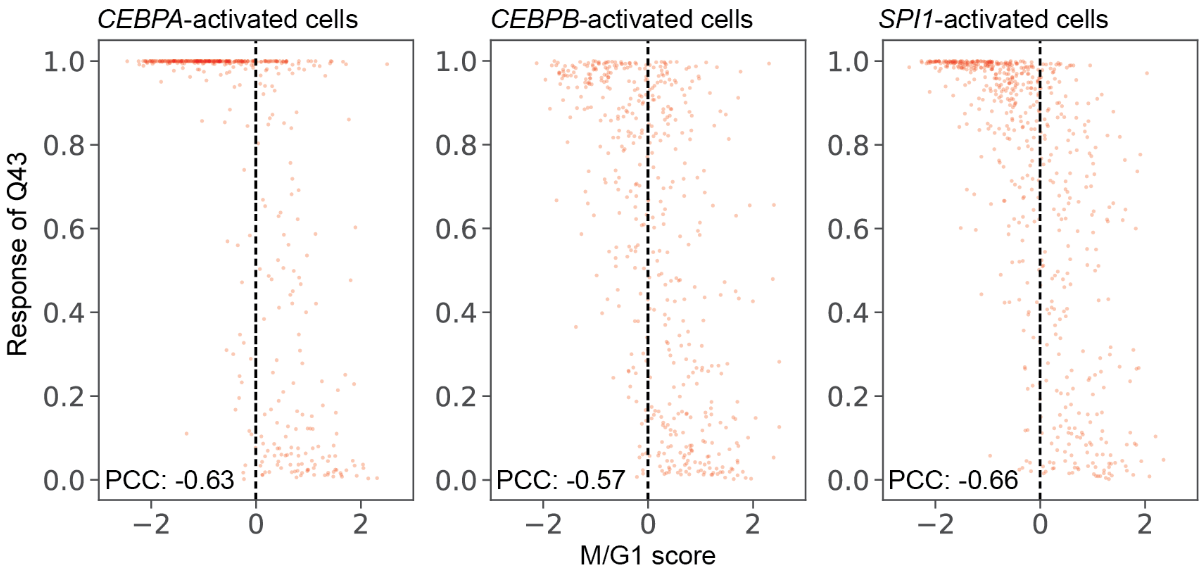


Figure S 17: Comparison of M/G1 score and response amplitude of Q43 in perturbations *CEBPA*, *CEBPB*, and *SPI1* from the K562 Perturb-seq data from Fig. 4.

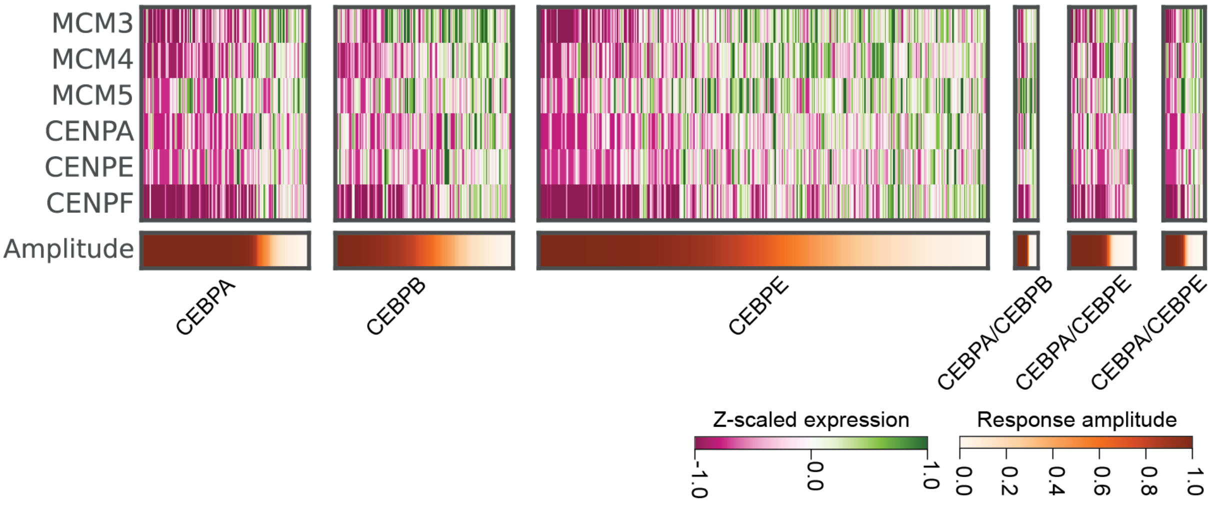


Figure S 18: K562 Perturb-seq data from Fig. 4. Expressions of *MCM3*, *MCM4*, *MCM5*, *CENPA*, *CENPE* and *CENPF* under conditions of CRISPR activation of *CEBPA*, *CEBPB*, *CEBPE*, *CEBPA/CEBPB*, *CEBPA/CEBPE*, and *CEBPB/CEBPE*. Cells are ordered based on response amplitudes of Q43.

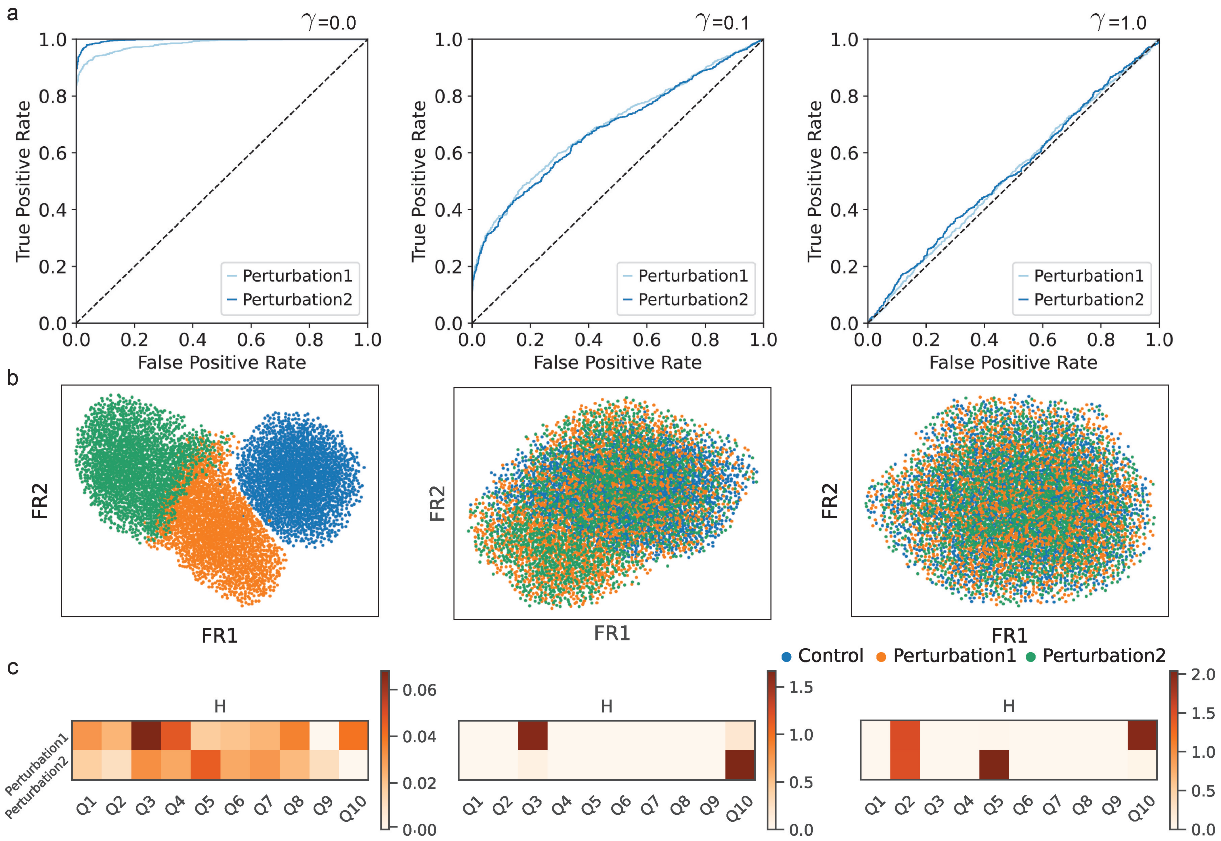


Figure S 19: Impact of the $\gamma$ hyperparameter (see Eqn. 19) on learning well-mixed basal state representations in simulation scenario 3. (a) ROC curves for the adversarial classifier. (b) Visualization of the inferred basal states in the Fruchterman-Reingold (FR) layout colored by experimental condition. (c) The inferred transcriptional program usage in each perturbation, $H_{pq}$. For small $\gamma = 0$, the basal state fails to achieve good mixing between the perturbation conditions. The model fails to learn response programs (note the scale on the color axis) since the basal state subsumes the perturbation responses. With $\gamma = 0.1$, we begin to see mixing of perturbation conditions in the basal state, but it is incomplete. With $\gamma = 1$, we achieve a basal state devoid of perturbation information, and we accurately recover the response programs.

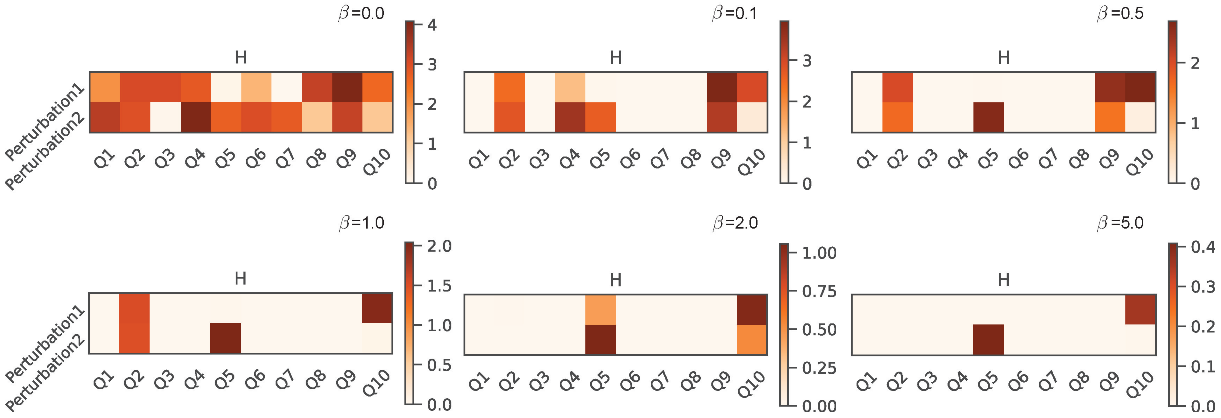


Figure S 20: Impact of the $\beta$ hyperparameter on inferring transcriptional program usage in each perturbation, $H_{pq}$, in simulation scenario 3. As we increase $\beta$, we increase the priority of automatic relevance determination in the loss function (Eqn. 19), and we see that CellCap finds fewer relevant response programs. $\beta$ values $< 1$ lead to the inference of an over-abundance of response programs: a form of over-fitting. $\beta = 1$ leads to inference of $H_{pq}$ that matches the ground truth. As we increase $\beta$ further, we find even fewer relevant response programs (and we get a poorer fit to the dataset).

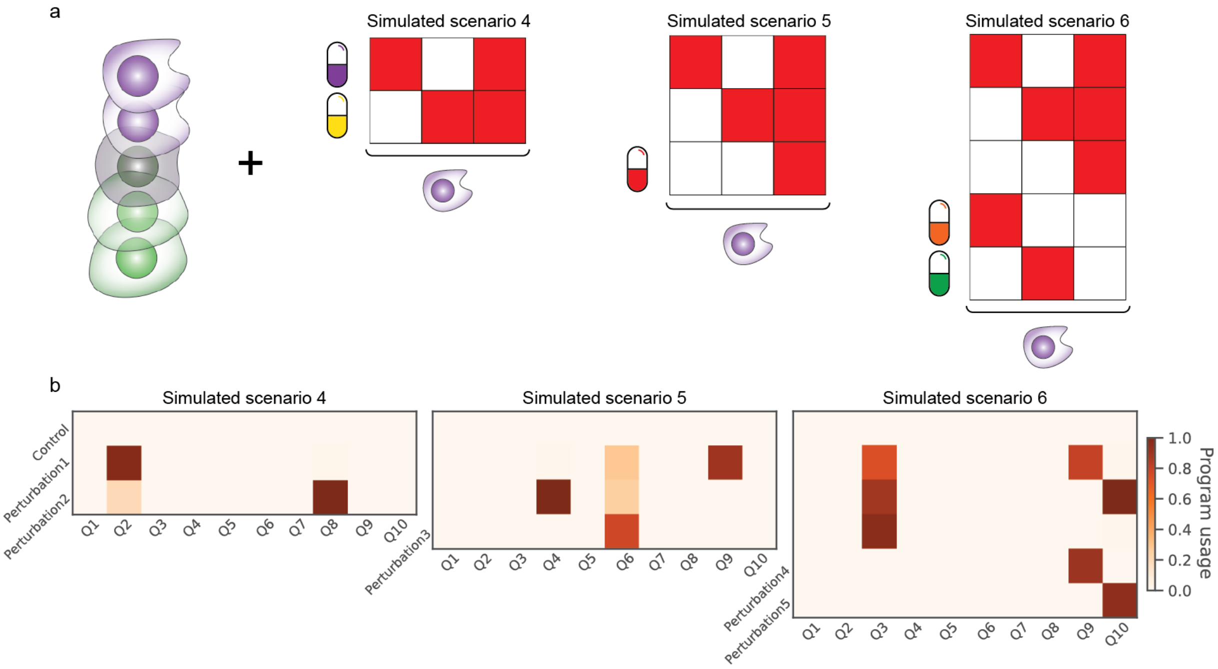


Figure S 21: CellCap results for the three extra simulated scenarios outlined in Supplementary Sec. S.1. (a) Graphical setups of the extra simulation scenarios. There are 3 transcriptional program in total as described in S.1. (b) The inferred usage of each transcriptional program for each perturbation in the three extra simulated scenarios.
